# Liver sinusoid constraints make random search of CD8 T cells for Plasmodium parasites efficient

**DOI:** 10.1101/2025.10.11.681610

**Authors:** Viktor S. Zenkov, Ian A. Cockburn, Raymond Lin, Whitney N. Griffith, Michael A. Langston, Vitaly V. Ganusov

## Abstract

A relatively small number of liver-localized CD8 T cells can provide sterilizing protection against exposure to Plasmodium sporozoites. Previously published mathematical model-based simulations of randomly searching for an infection T cells, assuming that T cells move in open (3D) space or on a regular lattice, suggested that some degree of attraction is needed for cells to find the infection. Yet, in our previous experiments we failed to find evidence of attraction towards the infection for most T cells. Because the movement of liver-localized T cells is confined to sinusoids, the blood vessels of the liver, T cells may in fact be attracted to the infection site via the sinusoids. In new experiments we imaged Plasmodium-specific CD8 T cells, sporozoites, and liver sinusoids simultaneously. We also developed a new method to model imaged sinusoids as graphs and to simulate movement of T cells on graphs, and new tests to detect biased cell movement on graphs. As before, our previously published open space/3D-based metric suggested that few T cells are attracted to the infection site in presence of moderate cluster of T cells. However, with our new graph-based metric we could not detect attraction when constraining T cell movement to sinusoids. Surprisingly, simulations of T cells searching for an infection via liver sinusoids showed that a relatively small number of T cells (1 cell/imaging volume or *<* 10^5^ cell/liver), moving without attraction at experimentally observed speeds, is sufficient to find the infection within 24 hours, well below the expected lifespan of the Plasmodium liver stage in mice. Our results suggest that the constrained environment of the tissue structure makes a random search more effective, and that detecting T cell attraction towards specific areas may arise as an artifact of metrics designed for open space/3D. Our results thus call for a re-evaluation of the implicit assumption that attraction is necessary for T cells to find infections in tissues.

## 1 Introduction

Malaria, a disease caused by eukaryotic parasites from the genus Plasmodium and transmitted by mosquitoes, remains unvanquished. Despite years of interventions such as bed nets, improved control of mosquito populations, and effective treatments, every year there are still millions of clinical cases of malaria and thousands of deaths, primarily in children under 5, from malaria [1–3]. The development and recent approval by the WHO of the two malaria vaccines, RTS,S and R21, is a milestone, but a relatively low efficacy and short duration of protection afforded by the vaccines suggest that more work is needed to control and eliminate malaria [4–7].

Mosquitoes cause malaria by inoculating a specific stage of malaria parasites, sporozoites (**SPZs**), into the skin during probing [8, 9]. Skin-deposited SPZs move and invade skin blood vessels, travel to the bloodstream, then invade hepatocytes and form liver stages [8]. It takes 2 days in mice (or about 6-7 days in humans) for each liver stage (**LS**) to develop into thousands of merozoites which then enter the circulation, infect red blood cells, and ultimately cause fulminant disease [8, 10–13]. RTS,S and R21 vaccines induce SPZ-specific antibodies (**Abs**) that neutralize the SPZs in the skin and/or blood, thus preventing clinical disease [7, 14–17].

Once a SPZ invades a hepatocyte, it is invisible to the action of Abs, and Plasmodium-specific CD8 T cells are required to clear infected hepatocytes [18–23]. Interestingly, in mice, CD8 T cells specific to a single epitope of murine Plasmodium berghei (**Pb**) or P. yoelii (**Py**) SPZs can provide sterilizing protection against relatively large infection doses [19–22, 24]. Protective CD8 T cells can be generated by vaccination or simply by transferring sufficiently large numbers of activated Plasmodium-specific CD8 T cells intravenously [19, 25–28].

However, how exactly T cells provide protection against SPZ challenge remains incompletely understood. To achieve sterilizing protection, T cells have *∼*48 hours to survey *∼* 10^8^ hepatocytes and eliminate all liver stages because full development of just one LS is sufficient to cause the blood stage disease [29]. Previous studies suggested that liver-localized (or liver-resident) CD8 T cells are required for protection, but recruitment of blood-circulating T cells to the liver after SPZ infection has been also suggested to play a role [26, 30]. We previously showed that T cells can form clusters around Plasmodium liver stages; such clusters are formed relatively rapidly (within a few hours after infection or T cell transfer) and the formation of the clusters was required to eliminate a liver stage [31, 32]. Our more recent analysis of the kinetics of LS elimination by T cells suggested that T cell clusters increase the probability of a liver stage’s death due to the highly variable killing abilities of individual T cells [33].

Factors determining formation of T cell clusters around Plasmodium liver stages remain poorly understood. We previously showed that the data on the distribution of T cell cluster sizes between different liver stages is best explained by the density-dependent recruitment (**DDR**) model whereby the first T cell searches for the parasite randomly and upon finding the target, it attracts other T cells to the infection site [31, 32]. However, by carefully tracking movements of T cells by intravital microscopy, we found that most T cells in the imaging volume did not display measurable attraction towards the liver stage; only in the case of a relatively large T cell cluster (5-7 cells) did we observe a bias towards the infection for a minority (*∼* 15 *−* 20%) of moving T cells [34]. The lack of detectable attraction for most imaged T cells was unexpected given that our and other agent-based simulations suggested that a random search of T cells for the liver stage would take too long (or would require too many T cells, *∼* 10^7^*/*liver) for sterilizing protection [26, 34].

In our previous work evaluating attraction of T cells towards an infection site, we made the common simplifying assumption that the direction of attraction should be from T cells directly towards the liver stage, i.e., assuming open space/3D environment [34]. However, we and others have previously shown that liver-localized CD8 T cells move primarily in liver sinusoids, specialized blood vessels of the liver [35], and therefore T cell attraction towards the infection site may not follow a direct Euclidean path [27, 36–38]. We and others have also shown that simulated T cells exhibiting correlated random walks based on existing open space-based methodology can be highly efficient at finding targets in constrained 1D environments [38, 39]. However, our previous study did not take actual liver structure into account in simulations, nor it established how the number of T cells in the liver determine the time it takes to find the infection especially in presence of bias of T cell movement towards the infection site [38].

Here, we extend our previous work and investigate if Plasmodium-specific CD8 T cells exhibit bias towards the parasite when we take the *real* structure of liver sinusoids and the constrained movement patterns of liver-localized CD8 T cells into account. Specifically, we performed experiments in which we used intravital microscopy to track locations of the Plasmodium liver stages and liver-localized CD8 T cells over time and imaged liver sinusoids in the same setting. We converted 3D images of liver sinusoids into graphs with vertices denoting branching points of the sinusoids and edge weights corresponding to the lengths of the sinusoids between vertices. We also developed methodology to simulate movement of T cells on these graphs with a variable degree of attraction or repulsion towards the liver stage (LS is defined as a collection of vertices) via a shortest path [40], and statistical methods to detect bias in the movement of T cells towards an infection site. Surprisingly, in our experiments we found similar numbers of T cells detected as attracted or repulsed via sinusoids, suggesting a lack of bias in moving towards the infection, even though a small proportion of T cells exhibited a bias towards a moderately sized T cell cluster in open space/3D. These results suggest that liver-localized T cells moving in sinusoids may search for the infection site randomly. Simulating random and unbiased T cell search for the infection site on our liver sinusoids-derived graphs revealed the high efficacy of such a search, with an average of 12 hours taken by a single CD8 T cell in the imaging volume (or about 6 *×* 10^4^ cells/liver) to find the parasite. Our results thus suggest that detecting any attraction of moving T cells towards an infection when using open space/3D attraction metrics may actually be incorrect unless specifics of the tissue structure used by T cells for movement is taken into account. Furthermore, an unbiased (no attraction) search may in fact be sufficiently efficient if the search occurs in biologically constrained tissues such as liver sinusoids and at reasonable speeds.

## 2 Materials and methods

### 2.1 Experimental design

#### 2.1.1 Mice

C57BL/6J (CD45.2) mice, B6 CD45.1, and OT-I mice [41] were purchased from the Jackson Laboratory and bred in-house at the Australian National University (**ANU**) under specific pathogen-free conditions except during infection experiments. Mice were age-matched between 6-8 weeks, and were sex matched for all experiments. All animal procedures were approved by the Animal Experimentation Ethics Committee of the ANU (Protocol numbers: 2019/36; A2022/36). All research involving animals was conducted in accordance with the National Health and Medical Research Council’s Australian Code for the Care and Use of Animals for Scientific Purposes and the Australian Capital Territory Animal Welfare Act 1992.

#### 2.1.2 In vitro activation of CD8 T lymphocytes

Single cell suspensions of C57BL/6 splenocytes were obtained from euthanised animals and incubated with 1*µ*g/ml of SIINFEKL ovalbumin peptide to stimulate T cells. The cells were then co-cultured with a single cell suspension of OT-1 splenocytes in T75 tissue culture flask (ThermoFisher) for 2 days. On day 3, cells were sub-passaged into fresh complete RPMI supplemented with 12.5U/ml of recombinant human IL-2 (rhIL-2, Peprotech) and incubated for a further 24 hours. The cells were sub-passaged a final time with fresh media and IL-2 before being purified on a Histopaque^®^ gradient and transferred.

#### 2.1.3 Adoptive transfer of CD8 T lymphocytes

OT-I cells were purified on a Histopaque gradient post activation in-vitro or eluted from a CD8-negative selection MACS column (Miltenyi) from single cell suspensions of splenocytes. Once purified, the cells were stained (CellTrace™ Violet/ CellTrace™ CFSE) diluted and transferred i.v into sex matched C57BL/6 recipients unless otherwise indicated (**Figure 1A**). For intravital imaging and lymphocyte tracking experiments, 7 *×* 10^6^ cells were transferred to each mouse.

**Figure 1:**
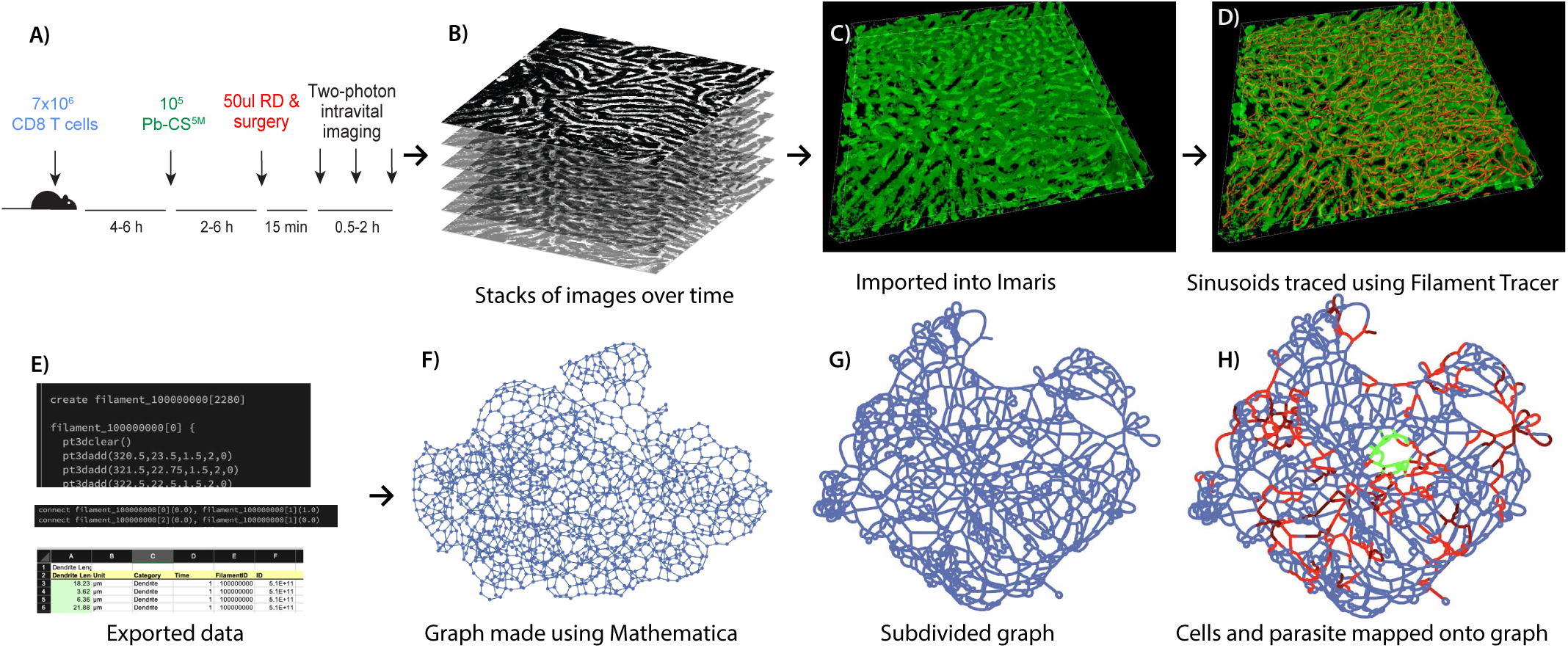
The experimental design and processing pipeline to generate a graph-based representation of liver sinusoids. We developed methods to image the sinusoid structure, model the structure using graphs, and model T cell movement in the sinusoids using graph algorithms. **A**: We injected 7 *×* 10^6^ in vitro activated OT-1 CD8 T cells, specific to Pb-CS^5*M*^, then 4-6 hours later we injected 10^5^ Pb-CS^5*M*^ sporozoites; after 2-6 hours we injected rhodamine dextran (**RD**), and prepared mice for imaging by exposing the liver 15 minutes later. **B**: We used a two-photon microscope to image a volume approximately 512 *×* 512 *×* 46 *µ*m^3^, collected as a z-stack of 2D images 512 *×* 512 *µ*m^2^ that are 2 *µ*m apart. We imaged the z-stack every 15 seconds to 2 minutes over a period of 30 minutes to 4 hours. **C**: We imported the z-stacks to the image analysis software Imaris. **D**: We used Imaris’ extension Filament Tracer to create filaments representing sinusoids. **E**: We exported details of filaments, traced using Imaris, into the .hoc file format. **F**: Using details of digitized sinusoids, we constructed a graph in Mathematica, with vertices corresponding to intersections of sinusoids and edges representing the length of the sinusoids (with the edge’s weight as the length of a sinusoid in *µ*m). **G**: To model T cell movement on the graph, we *subdivided* graph edges with weights *>* 3 *µ*m; this added around 10,000 vertices with degree 2 to the graph (**Table 1**). **H**: We digitized positions of CD8 T cells (shown in red) and of the parasite (shown in green) and mapped them to vertices on the graph. Because sporozoites reside inside of hepatocytes, we assumed that the infected hepatocyte is a 40 *µ*m sphere [32] and considered every vertex inside that sphere to be an infected vertex, with around 100-400 vertices per liver stage (**Table 1**).

**Table 1:** Details of individual experiments of imaging liver-localized CD8 T cells, Plasmodium sporozoites, and liver sinusoids, and characteristics of graphs based on the digitized sinusoid structure. We re-analyzed imaging data from previous experiments and performed new experiments in which we tracked movement of liver-localized CD8 T cells while labeling liver sinusoids; in total we have 5 datasets dubbed MouseC, MouseD, CTV3, CTV5, and Mouse4. Datasets are primarily distinguished based on the size of the cluster (the number of T cells within 40 *µ*m of the parasite), while the MouseC and MouseD experiments used mice that were uninfected and are from our previous publication [38]. Note that the duration of the CTV3 experiment is short because the mouse shifted dramatically partway through imaging, so only a portion of the experiment could be used. For each experiment, we generated a graph, with edges representing sinusoids and vertices representing intersections of sinusoids, and edges’ weights set to the length between two branching points of a sinusoid (*µ*m). We also generated a *subdivided* graph, where we subdivided each edge with weight over 3 *µ*m into multiple shorter edges with equal weights below 3 *µ*m (**Supplemental Figure S1**). The infected hepatocyte is modeled as a 40 *µ*m sphere surrounding the parasite, which encompasses multiple (“infected”) vertices of the graph. For each graph and subdivided graph, we provide the following graph characteristics: the numbers of vertices and edges, the density, the radius (the minimum eccentricity, i.e. the smallest distance from a vertex to any other vertex), the global clustering coefficient (**c.c.**), and the fractal dimension (see main text for details of these graph metrics). We also compared the composition of these graphs by calculating distribution of local clustering coefficients (**Supplemental Table S1**).

| Experiment<br>Clustering details | Mouse C<br>No parasite | Mouse D<br>No parasite | CTV3<br>No cluster | CTV5<br>Small cluster | Mouse4<br>Supercluster |
| --- | --- | --- | --- | --- | --- |
| T cell cluster size | N/A | N/A | 0-1 | 2-3 | 10 |
| #sporozoites injected | 0 | 0 | $10^5$ | $10^5$ | $10^5$ |
| #T cells imaged | 88 | 124 | 70 | 87 | 61 |
| duration (min) | 46.97 | 64.61 | 13.42* | 31.12 | 26.20 |
| imaging frequency (min) | 0.401 | 0.422 | 0.497 | 0.502 | 0.364 |
| #vertices | 1457 | 2011 | 2497 | 1401 | 1171 |
| #edges | 2175 | 3402 | 4116 | 2068 | 1719 |
| density | 0.0021 | 0.0017 | 0.0013 | 0.0020 | 0.0025 |
| #infected vertices | N/A | N/A | 41 | 23 | 26 |
| radius | 599 | 530 | 589 | 500 | 546 |
| global c.c. | 0.012 | 0.0061 | 0.0040 | 0.011 | 0.0097 |
| dimension | 2.07 | 2.11 | 2.15 | 2.12 | 2.01 |
| subdivided #vertices | 13862 | 11482 | 14716 | 13599 | 10387 |
| subdivided #edges | 14580 | 12873 | 16335 | 14266 | 10935 |
| subdivided mean degree | 2.104 | 2.242 | 2.220 | 2.098 | 2.106 |
| subdivided density | 0.00015 | 0.00020 | 0.00015 | 0.00015 | 0.00020 |
| subdivided #infected vertices | N/A | N/A | 323 | 198 | 188 |
| subdivided radius | 40568 | 33567 | 42709 | 39645 | 30233 |
| subdivided global c.c. | 0.00012 | 0.00095 | 0.00032 | 0. | 0.00016 |
| subdivided dimension | 1.66 | 1.65 | 1.68 | 1.71 | 1.69 |

#### 2.1.4 Infection of mice with Plasmodium sporozoites

Mice were infected intravenously (i.v.) via tail vein injection with 5 *×* 10^4^ *−* 10^5^ P. berghei CS^5M^ (Pb-CS^5M^) sporozoites, **Figure 1A** and [42]). Sporozoites were dissected by hand from the salivary glands of Anopheles stephensi mosquitos generated in-house within a quarantine approved facility.

#### 2.1.5 Labeling of liver sinusoids

To label the sinusoids, we injected Evans blue or Rhodamine dextran (**RD**) i.v. into mice (50 *µ*g/mouse) about 15 min prior to imaging of the livers (**Figure 1A**).

#### 2.1.6 Multiphoton microscopy

Mice were prepared for microscopy in vivo as described previously [43]. Once the mouse was ready and applied to the movable platform of a Fluoview FVMPE-R multiphoton microscope, the platform was raised to ensure contact of the XLPLN25XWMP2 objective lens with a drop of water on the coverslip (25x, NA1.05, water immersion; 2 mm working distance). For the analysis of motility of cells activated in vitro, an approximately 50 *µ*m Z-stack (2 *µ*m/slice) was typically acquired using the galvo-scanner at a frame rate of typically 2 frames per minute. The images were acquired using the FV30 software (Olympus) and exported to Imaris (Bitplane) for track analysis using the autoregressive motion algorithm and polarity analysis [38, **Figure 1A**].

### 2.2 Processing imaging data with Imaris

In this project we generated three novel movies and analyzed 5 datasets that include data from our previous publication (**Table 1**). All datasets featured T cells and sinusoids imaged as described above; the datasets labeled MouseC and MouseD were from uninfected mice, while the remaining datasets (CTV3, CTV5, and Mouse4) were from sporozoite-infected mice. The latter three datasets were distinguished by their “cluster size”, i.e. the number of T cells “near” the infection site: CTV3 represented the dataset with no cluster (0-1 T cells near the infection site), CTV5 represented the dataset with a small cluster (2-3 T cells near the infection site), and Mouse4 represented the dataset with a “supercluster” (10 T cells near the infection site, **Table 1**). In all cases, our data consisted of stacks of images of dimensions 512 *×* 512 *×* 46 *µ*m over time (**Figure 1B**) that we converted to Imaris’ format (.ims, **Figure 1C**).

#### 2.2.1 Determining position of the sporozoite

For each movie we recorded the position of the SPZ during acquisition of the data, as the sporozoite was clearly visible in the microscope. We then used the Spots function in Imaris using the green channel (SPZs express GFP) to determine the position of the sporozoite over time. While the SPZ was located inside of a hepatocyte and was thus nominally immobile, there was some minimal change in the parasite’s position over time due to the shifting of the liver in a breathing animal and/or due to movement of parasitophorous vacuole within the hepatocyte.

#### 2.2.2 Determining positions of T cells

We also traced positions of T cells using the Spots function in Imaris. This output a position in 3D for every time at which every cell was detectable in the imaging volume. In addition to automatic object tracking, we manually adjusted positions for every T cell to ensure that every position was accurate and that no time points were skipped for any cell - every cell’s positions were therefore consistently recorded from its first to its last moment present in the imaging volume. Manual tracing was also necessary to pick up cells that detached from the sinusoid walls and floated in the bloodstream at higher speeds, although such events are rare and do not affect our results (but see Discussion). Examples of Imaris parameters used for tracing the sporozoite and T cells are available in the Supplement.

#### 2.2.3 Quantifying sinusoids

Sinusoids are blood vessels in the liver [35], with a sinusoid precisely defined as the portion of the vessel between two branch points (places where there are three or more possible directions). We traced sinusoids using Filament Tracer, an extension of Imaris (**Figure 1D**). Filament Tracer can be applied to still images or movies; in the case of movies, filaments can be created for the entire imaging volume at each time point independently, which consumes much computer time and occasionally caused Imaris to crash. Furthermore, some variability in labeling of the sinusoids was expected due to flow of RD with the blood, and such variability did influence inference of the network of the sinusoids. However, since the structure of the sinusoids should not truly change over the imaging period of 1-4 hours, we typically chose a single representative frame and only traced sinusoids at that time point. We also inspected the automatically generated filaments and adjusted filament tracings manually. Parameters used in Filament Tracer are available with the Imaris file provided along with the coordinate data. We exported the object generated by the Filament Tracer into a special .hoc file format, which contains information on all branching points and connections between them (e.g., length between branching points, **Figure 1E**). We provide instructions for how to export data in the .hoc format in the Supplement.

#### 2.2.4 Correcting tissue drift using reference frames in Imaris

Because we performed intravital imaging of livers of live animals, we often observed shifting of the imaging volume, perhaps due to breathing or other movements of the animal. These tissue movements resulted in the imaging volume shifting over time by up to a few dozen micrometers, which is enough to change the locations of imaged cells significantly. It was thus necessary to adjust for this tissue movement. We used Reference Frames, another object in Imaris, that allowed us to synchronize shifts of the imaging volume between different frames. Coordinates of other objects such as the sporozoite, T cells, and of sinusoids are then shifted with respect to the reference frame coordinates.

### 2.3 Representing sinusoids as graphs

We next transformed Imaris’ tracings of sinusoid structures into *graphs*. (See the Supplement for technical information defining graphs.) For each experiment, the output of this transformation was a list of *vertices* and a list of *edges*, with each edge accompanied by a numerical *edge weight*. We define the graph as follows:

1. Vertices are set to intersections of sinusoids (defined in the .hoc file).
2. Edges are set to sinusoids (defined in the .hoc file).
3. An edge weight is set to the length of the edge’s corresponding sinusoid (attained by summing the distances between subsequent points in a traced filament).

As previously stated, the graph was constructed for the single frame we chose to trace in Filament Tracer, so all other aspects of the movie are adjusted with respect to that time point. The vast majority of vertices of every graph have degree 3 (**Figure 1F** and **Supplemental Figure S1**), as every intersection of sinusoids by definition has at least three “options” of paths for a cell to travel. A minority of vertices have degree 4 or 5, and some vertices near the boundaries of the imaging volume may artificially have degree 2.

A liver sinusoids-derived graph is biologically expected to be “connected”, i.e. there is a path from any vertex to any other vertex, because blood vessels of the liver are fully connected. However, since the imaging volume is only a sliver of the liver, it is possible that small numbers of vertices only connect to the remaining vertices via sinusoids outside the imaging volume and there are thus separate “components” of the graph. These vertices may be ignored and do not affect our analyses.

In graph theory, there is no concept of being “at a location along an edge” - individual vertices are the only “locations” which have any meaning. Since real cells are not restricted to intersections of sinusoids (the vertices in our graph), and indeed often move a smaller distance between each recorded position in the experiment than the length of sinusoids (the edges), cells are very commonly situated on sinusoid branches and not at or near intersections. Since each cell position must correspond to a vertex to be usable in a graph-based framework, we split many edges of the graph, a process called *subdivision*. In particular, we subdivided every edge of the graph into a chain of edges with equal edge weight (length) such that each edge weight was less than 3*µ*m. This increased the number of vertices and number of edges by the same quantity, adding around 10,000 vertices with degree 2 (**Figure 1G**).

### 2.4 Calculating fractional dimension of the graphs

There exist several metrics estimating fractal dimension. One commonly used is “box dimension” [44], which is not effective on graphs [45]. We use a definition specifically designed for graphs [46]. The estimate for dimension is

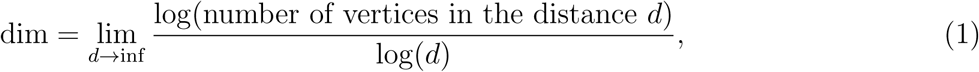

where *d* is a distance (accumulated weight) from a chosen starting vertex. In words, this is the limit of the relationship between a traversal distance and the number of vertices visited at that distance.

This algorithm breaks down as the traversal reaches boundaries of a graph, as the number of new destinations is artificially cut off. This is especially consequential for our sinusoid graphs, created from thin 500 *×* 500 *×* 50 *µ*m^2^ slices of the liver. To alleviate this, we extrapolate the graph from its 500 *×* 500 *µ*m^2^ faces by repeatedly duplicating the entire graph and connecting vertices at the boundary (defined as being within 3 *µ*m of the maximum value in one axis) in the original graph to the duplicates of those vertices in the new graph. By repeating this process 9 times, we have a graph representing a 500 *×* 500 *×* 500 *µ*m^3^ cube consisting of 10 graphs representing the same imaging volume stacked upon each other.

### 2.5 Mapping positions of T cells onto the graph

To use cells in the context of our liver sinusoids-derived graphs, we mapped the positions of every cell to the vertex on the subdivided graph whose corresponding real position was closest to the recorded T cell position via Euclidean distance. The result is a list of vertices visited by each cell at subsequent times after the start of imaging.

#### 2.5.1 Defining vertices visited by T cells

Given the average speed of liver-localized CD8 T cells of 5 *−* 15 *µ*m*/*min (with occasional faster displacement with blood flow) [27, 34, 38], it was possible for a given T cell to move more than 3 *µ*m between two sequential time frames. For such movements, a cell moved more than one vertex on the subdivided graph. In our estimation of whether a cell moves closer to the parasite, we need to track all vertices that a cell visited in one movement. We therefore calculated the shortest path on the graph (i.e. the list of consecutive adjacent vertices with the shortest total edge weight) between each successive pair of vertices visited by cells to be a hypothesis of the path taken by the cell between each recorded position. We colloquially refer to the steps between time steps as “midsteps”. Because T cells in the liver rarely turn around [38], these hypothesized paths should be relatively accurate. However, it is not possible for us to know exactly what path is taken between recorded positions if a T cell moves by multiple vertices between two sequential time frames. In contrast to these lengthy tracks, if a cell does not move in one time period, it will still map to the same vertex, so small movements are absorbed because vertices are not continuous.

#### 2.5.2 Mapping the parasite’s position to multiple vertices

We also mapped the parasite onto the graph in a similar manner to the T cells. We mapped the first recorded parasite position to its nearest vertex and considered that to be the “main” infected vertex. Murine hepatocytes vary in size, and following our previous calculations, we estimated a hepatocyte as a sphere with 40 *µ*m radius [32]. All vertices that are located within this distance from the “main” infected vertex are therefore considered to be part of the infected hepatocyte. In the subdivided graph, out of *∼*10,000 total vertices, about 100-400 vertices were labeled as “infected” (**Table 1**).

#### 2.5.3 Representing boundaries of the imaging volume

We considered every vertex within 3 *µ*m of a boundary of the imaging volume (where the boundaries are the minimum or maximum values in each coordinate direction) to be a *boundary vertex*. Such vertices’ corresponding positions on the sinusoid structure may connect to sinusoids that traverse outside of the imaging volume, which by definition are not imaged or represented in the graph.

### 2.6 Defining distance between a T cell and the parasite via shortest path weights (SPWs)

For a given time point, there are multiple paths between the vertex where a given T cell is and the set of “infected” vertices representing the parasite. Along each such path we can calculate the path weight that is the total sum of edge weights along the path, which is the distance from the cell to the parasite within the structure. We also calculated the *shortest path weight* (**SPW**), which is the total weight of the shortest path between one vertex (T cell’s location) and a collection of “infected” vertices (parasite). We calculated the SPW from each vertex in the graph to each “infected” vertex in the graph using a method based on Dijkstra’s algorithm [40], and then chose the smallest SPW to be the SPW from the T cell to the parasite.

### 2.7 Metrics to test for attracted movement

To evaluate whether moving T cells display attraction towards the infection site, we use several metrics. One metric is based on the Von Mises-Fisher (**VMF**) distribution, which we previously found to be the most sensitive to detect attraction in open space/3D settings [34]. To detect potential T cell attraction towards the infection site via sinusoids, we developed novel metrics for estimating attraction on graphs.

#### 2.7.1 Von Mises-Fisher (VMF) distribution-based metric for 3D/open space

To quantify attraction (or repulsion) of cells towards an infection site, for every cell movement we calculate the angle *φ* between the movement vector and the vector towards the infection site. For cells moving without attraction, the distribution of these angles to the infection is given by sin(*φ*). Deviations of random movement (with respect to the infection) in 3D can be evaluated by using the VMF distribution:

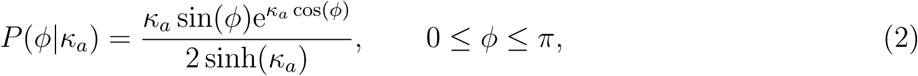

characterized by the concentration parameter *κ_a_*. The concentration parameter indicates the “strength” of attraction (*κ_a_ >* 0) or repulsion (*κ_a_ <* 0) [34]. To estimate *κ_a_* for a given T cell or all T cells in the movie, we calculate all angles to infection *φ_i_* and use a maximum likelihood method:

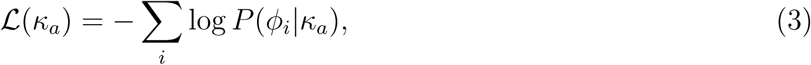

where *L* is the negative log-likelihood. To evaluate if *κ_a_* is statistically different from 0, we use a likelihood ratio test (**LRT**) to compare the negative log-likelihood of the best fit (*L*(*κ̄_a_*)) and the negative log-likelihood with lim*_κa→_*_0_ *L*(*κ_a_*). The quality of the VMF distribution fit to the angles to infection can be also evaluated visually (e.g., **Supplemental Figure S8**). Of note, we previously showed that the VMF distribution-based method is more sensitive at detecting attraction than methods based on average angles to infection or on classifying movements as being “towards infection” or “away from infection” [34].

#### 2.7.2 SPW-based metric to detect attraction on graphs

Similar to the VMF distribution-based metric, we define an attraction parameter *α* that determines the probability of a cell moving from vertex *i* to vertex *j* based on the change in SPWs for these vertices as

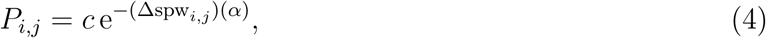

where Δspw*_i,j_* = spw*_j_ −*spw*_i_* and *c* is the normalization constant that is calculated by summing over all potential paths a T cell can make in one time frame (**Figure 2**). For example, for a cell moving from vertex 0 with a SPW of 3 to vertex 1 with a SPW of 1, the change in SPW is Δspw_0_,_1_ = 1 *−* 3 = *−*2 and the probability of the movement is then *P*_0_,_1_ = *c* e^2^*^α^* (**Figure 2Bi**). The probability of staying in the same vertex (**Figure 2Bii**) or moving farther from the infection site (**Figure 2Biii**) is calculated similarly.

**Figure 2:**
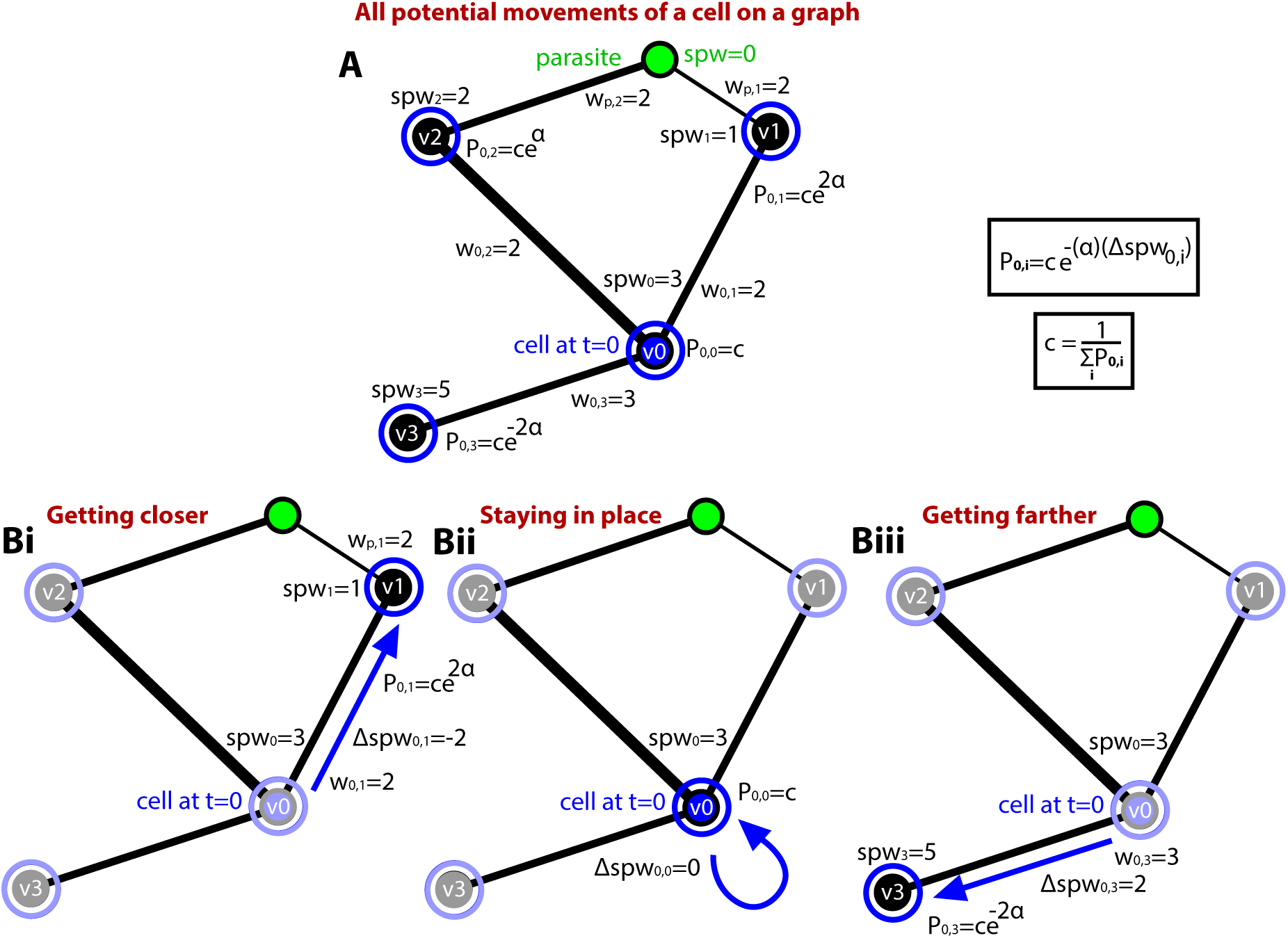
Methodology to model movement of a cell on a graph with a bias towards an infected vertex. We mapped digitized positions of T cells and the parasite to specific vertices of the graph and defined probabilities of T cell movement between vertices, with bias in movement towards an infected vertex defined by a parameter *α* (**eqn. (4)**). While a cell may move a distance corresponding to many vertices in a single timestep, that cell does visit a list of adjacent vertices across its midsteps, so every time a cell is to move it will move to a vertex adjacent to the current vertex - or to the current vertex itself (staying in place for the entire timestep). **A**: Initially, a cell is located at vertex *v*_0_ and may stay in place or move to vertices *v*_1_, *v*_2_, or *v*_3_ with the probability *P*_0*,i*_ = *c* e*^−α^*^Δspw0^*^,i^*, where Δspw_0*,i*_ = spw*_i_ −* spw_0_ and spw_0_ is the shortest path weight (**SPW**) from the current cell’s vertex to the parasite’s vertex and *c* is normalization constant. **Bi**: The probability of moving from a vertex *v*_0_ to vertex *v*_1_ is *P*_0,1_ = *c* e^2*α*^, given the change in distance Δspw_0,1_ = *−*2. **Bii**: The probability of staying at the same vertex *v*_0_ is *P*_0,0_ = *c* since Δspw_0,0_ = 0. **Biii**: The probability of moving from vertex *v*_0_ to vertex *v*_3_ is *P*_0,3_ = *c* e*^−^*^2*α*^ since Δspw_0,3_ = 2. In simulations, we pseudrorandomly choose the movement of a cell given weighted probabilities of each displacement. When cell displacement per time frame is larger that any of change in SPW we divide the movement into midsteps (**eqn. (9)**) and simulate cell movement as in panel B for each of the midsteps (see Materials and methods for details).

We calculate the probability that the cell gets closer to the parasite (Δspw*_i,j_ <* 0) out of all possible movements for every movement at time *t*:

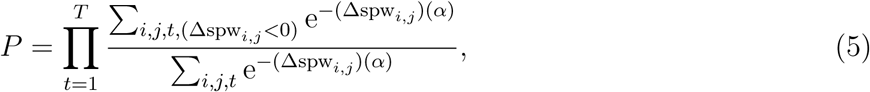

where *T* is the number of time frames in the movie. We then calculate the negative log-likelihood as

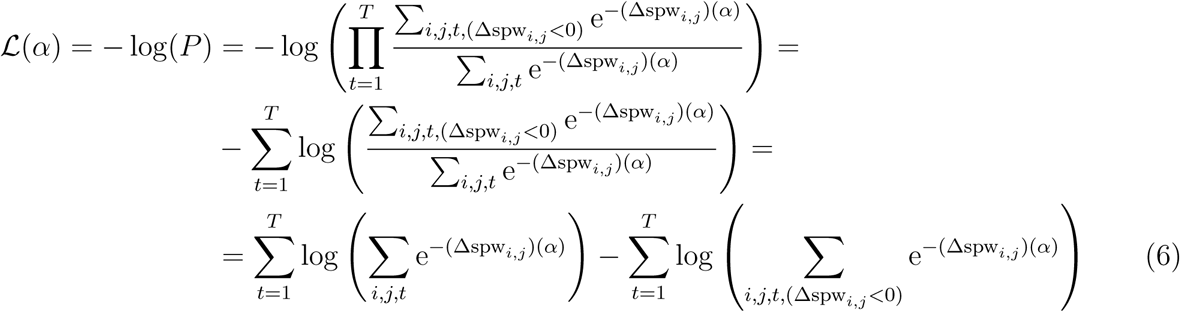

where we find the best fit *α* by minimizing *L*. To evaluate if *α* is statistically different from 0, we use a LRT where we compare *L*(*α*) with lim*_α→_*_0_ *L*(*α*) [47].

#### 2.7.3 Alternative metrics to detect attraction on graphs

We also formulated and tested several other metrics to detect attraction of T cells towards the infection site. In the **monotonicity** metric, we calculated the SPW for each T cell position and calculated the Spearman rank correlation coefficient between SPW and time. A statistically significant decline in SPW with time may indicate that a T cell is “attracted” to the infection site. Alternatively, we compared the average of all SPWs for all cell movements to the initial shortest path weight by using a **t-test**. If the average SPW has significantly decreased, then the cell is classified as attracted.

#### 2.7.4 Statistical methodology to define attracted cells in a collection of cells

We classify each T cell as unbiased or biased (where bias encompasses attracted or repulsed) with any of our metrics (e.g., **eqn. (2)** or **eqn. (5)**). To determine if a *collection* of cells are attracted to the infection site, we calculate the fraction of attracted cells among biased cells as

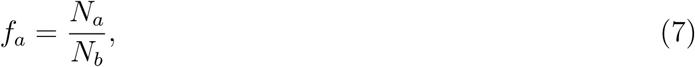

where *N_a_*= number of attracted cells and *N_b_*= number of biased cells (*N_a_*+ *N_r_*, where *N_r_* is the number of repulsed cells). Bias in attraction is then calculated using the binomial distribution with parameters *p* = 0.5 and *n* = *N_b_*, and *N_a_* as the number of successes [34].

We note that the binomial test cannot be used without taking into account quantities of cells: if, for example, there were 10,000 T cells, and only 20 were biased, but all 20 were attracted, the binomial test would describe the collection of cells as attracted, despite the vast majority having no bias. In this light, we developed the **combo** test, which instead tests the fraction

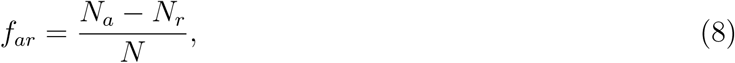

where *N* is the total number of cells and we test *f_ar_* using a binomial test with *p* = 0 and *N*.

### 2.8 Simulating T cell walks on graphs with degree of motility and bias toward the parasite

Given a liver sinusoids-derived graph, with some vertices defined as “infected”, we simulated movement of T cells using two main parameters: the attraction parameter *α* and a motility parameter *ψ*. Distances moved per time frame are sampled from the displacements of actual T cells from a given experiment/movie.

We position a cell at a pseudorandomly chosen starting vertex. At every timestep, the cell will either remain at its current vertex or make a move of approximate distance *d* sampled from the distribution of distances moved between vertices. That distance *d* is divided into *m* steps, one per midstep, where *m* = *[m_c_|* is the smallest integer greater than

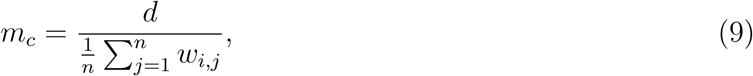

where *w_i,j_* is the edge weight connecting vertices *i* and *j*, and 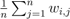 is the average edge weight for all *n* edges connected to the current vertex *i*. The probability that a cell moves from vertex *i* to vertex *j* is given by

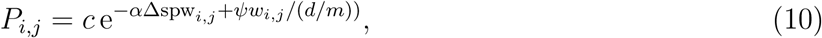

which is extended from **eqn. (4)** by adding a probability of a cell staying or leaving the current location, dependent on *ψ*, where *c* is the normalization constant. We then select the movement of a cell according to normalized movement probabilities from **eqn. (10)**. If the chosen vertex is the current vertex, the process ends; otherwise the cell moves to the chosen vertex and the process is repeated for the adjacent vertices (now not including the prior vertex or the current vertex, as neither turning-back nor staying-in-place can be measured between timesteps of real cells), and the cell continues to move for *m* midsteps total.

Cells thereby have attraction (defined by *α*) and motility (defined by *ψ*), and both quantities may be negative (indicating repulsion or a preference to move away from the current vertex, respectively). We simulated the movement of T cells on sinusoids-derived graphs and by comparing the proportion of movements that stay in place in data and simulations, we found the value of *ψ* to mimic real cells’ movements to be about *ψ* = *−*1.2 in movies/datasets CTV3, CTV5, and Mouse4 (SPZ-infected mice) and *ψ* = *−*1.45 in movies/datasets Mouse C and Mouse D (uninfected mice). We chose values of the attraction coefficient *α* to vary from negative (repulsion) to positive (attraction) values, with *α* = 0 as unbiased cell movement. To reduce the number of vertices that a cell visits in one time frame, we set the limit of one displacement to 100 *µ*m. This limit contributes slightly to simulated cells moving on average slower than real cells. Given that our movies were acquired with a timestep of about 30 sec, our assumed speed limit of 200 *µ*m*/*min likely includes both crawling and floating-in-blood T cell movements [38, 48, 49]. Large displacement events typically constitute a small proportion of all movements (e.g., **Supplemental Figure S5**).

For simulations of T cell movement for 48 hours, the time limit to find and kill parasites to stop malaria infection in mice, we modeled the ability of a cell to leave the imaging volume. When the cell is on a boundary vertex (defined above), it has a probability of “leaving” the imaging volume if it remains on a boundary vertex for two time steps. To keep the number of cells searching for the parasite in the imaging volume constant, when a cell left the imaging volume, we allowed a new cell to “enter” the imaging volume at a psuedorandomly selected boundary vertex.

### 2.9 Additional computational details

We performed our stochastic simulations in Mathematica 13.1.0. To run simulations of T cell movements for 48 hours we used parallelized code on the ISAAC supercomputer of the University of Tennessee (https://nics.utk.edu/isaac/).

## 3 Results

### Imaging T cells, sporozoites, and liver sinusoids in one setting and converting images of liver sinusoids to graphs

Over the years we have developed the expertise to image Plasmodium sporozoites (**SPZs**) and liver-localized CD8 T cells in livers of live, anesthetized mice using spinning disk or two photon microscopy [16, 27, 31, 43]. In such experiments we typically use SPZs expressing GFP and label T cells with fluorescent labels in vitro prior to their adoptive transfer and imaging. In other recent experiments we also developed a robust methodology to label liver sinusoids by injecting rhodamine dextran (**RD**) 10-20 min prior to imaging; with this methodology we recently quantified movement programs of liver-localized CD8 T cells and identified how SPZ-specific antibodies (**Abs**) influence the ability of individual parasites to invade hepatocytes [16, 38]. Here we extended our previous studies to image SPZ, liver-localized SPZ-specific CD8 T cells, and liver sinusoids in one setting (**Figure 1**). Specifically, we first transferred 7 *×* 10^5^ in vitro activated OT-I CD8 T cells, specific to SIINFEKL peptide from chicken ovalbumin. Then 4-6 hours later we injected i.v. 10^5^ GFP-expressing *Plasmodium berghei* (**Pb**) SPZs expressing SIINFEKL peptide in circumsporozoite protein (Pb-CS^5M^). After another 2-6 hours we injected i.v., RD and 15 min later, exposed and imaged the liver with a two-photon microscope (**Figure 1A** and **Movie 1**).

In the scanned volume (**Figure 1B**) we identified locations of the SPZ within hepatocytes in the microscope and then used the Spots object in Imaris to track parasite’s position over time; a parasite’s coordinates may shift due to movements within the parasitophorous vacuole within the hepatocyte and/or because of tissue shifts due to mouse breathing (**Movie 2**). Similarly, we used Spots to track positions of adoptively transferred CD8 T cells in the liver over time (see Materials and methods for more detail and **Movie 3**).

We then used the Filament Tracer module in Imaris to track and quantify liver sinusoids (labeled with RD, **Figure 1C&D**). Importantly, Filament Tracer allows the creation of filaments covering all labeled blood vessels for the imaging volume and each time frame; however, we discovered that for longer movies Imaris was typically unable to generate filaments for each time frame (and would crash). Therefore, for each of our movies, we generated filaments of the imaged volume for a representative time frame; we used automatic tracing with manual editing of the generated filaments (e.g., **Figure 1D** and **Movie 4**). Imaris provides critical information about every filament (or “dendrite”) such as coordinates along each dendrite, a list of intersections of beginnings and endpoints of pairs of dendrites, etc that are exported in a specialized.hoc file format (**Figure 1E**; see Materials and methods and Supplemental Text for more detail). We used that information to convert generated filaments into graphs (or networks) by assigning intersections between filaments/dendrites as vertices (or nodes) and connecting vertices by edges as defined by each filament. Because the distances between different intersections of filaments vary, we assigned weights, corresponding to the sinusoid length between two intersections, to the edges connecting vertices (**Figure 1F**). Importantly, while general graphs are mathematical constructs and may not have direct relationship with the natural world, our liver sinusoids-derived graphs are directly related to the 3D structure of the liver. Therefore, every vertex in our graphs has associated 3D coordinates, allowing us to calculate the displacements of cells moving between vertices.

In total we generated graphs from five independent movies where we tracked the movement of liver-localized CD8 T cells in RD-labeled sinusoids (**Table 1** and **Supplemental Figure S1**): three movies generated in this study and two movies from our previous work [38]. Liver sinusoids-derived graphs had between 1,100 to 2,500 vertices and between 2,000 to 4,000 edges making these graphs relatively sparse with low density (*∼* 0.1%). In comparison, a typical density of graphs constructed from yeast gene expression data is 10 times higher, around 1% [50]. Low values of the global clustering coefficient of the liver sinusoids-based graph are also consistent with their sparsity (**Table 1**) [51]. The radii of these graphs are relatively large (*∼* 500) and consistent with the extended biological structure of the sinusoids (e.g., **Figure 1C**). Interestingly, the fractal dimension of these novel graphs was relatively small, around 2 [52], suggesting that in contrast with the overall 3D shape of the liver, the dimension of liver sinusoids is substantially smaller. This is consistent with the small contribution of the volume occupied by the liver sinusoids to the imaging volume of the liver in our movies (**Supplemental Figure S2**) and generally low degrees of freedom (indicated by low degrees of vertices). Overall, these graphs were similar in composition as judged by the distribution of local clustering coefficients (**Supplemental Table S1**).

To analyze movements of imaged CD8 T cells with respect to the parasite and liver sinusoids we next mapped the coordinates of T cells and an infected hepatocyte to vertices on the graph. Because in experiments we only track the position of the parasite but not the infected hepatocyte per se, we assumed that a hepatocyte is a sphere with a radium of 40 *µ*m [31, 32] and labeled any vertex within 40 *µ*m of the parasite’s position as “infected”. In the graphs, around 1.6-2.2% of the vertices (e.g., 23/1401) were defined as infected vertices (**Table 1**). While we and others have previously established that liver-localized CD8 T cells move primarily in liver sinusoids [36, 38], mapping positions of T cells to vertices of liver sinusoids-derived graphs proved somewhat difficult because some T cell positions appeared to be relatively far from 3D coordinates of vertices representing intersections of sinusoids. Therefore, to more accurately map 3D T cell positions to the graph vertices we subdivided the graph by creating additional vertices in edges that had weights larger than 3 *µ*m so that the maximum weight in our subdivided graphs would not exceed 3 *µ*m (**Figure 1G** and see Materials and methods for detail). Given the diameter of liver-localized CD8 T cells of about 10 *µ*m, such graph division ensures that for any 3D location, a T cell can be well mapped to a nearby vertex (**Supplemental Figure S3**). Such a mapping thus us allows to rigorously track the movement of T cells relative to the “infected” vertices over time (**Figure 1H**). Importantly, subdividing a graph naturally resulted in a smaller graph density, smaller global clustering coefficient, and substantially smaller dimension (*∼* 1.7, **Table 1**). On the other hand, more vertices in the subdivided graphs are defined as “infected”, and subdivision resulted in a much larger radius of each graph (**Table 1**). Taken together, for the first time we have developed a rigorous pipeline to convert 3D images of liver sinusoids into graphs and map movement patterns of liver-localized T cells and positions of Plasmodium liver stages to vertices on such graphs (**Figure 1**).

### Simulating and testing biased random walks on graphs

As we developed methodology to convert images of liver sinusoids into graphs and to map positions of T cells and the Plasmodium liver stage to vertices on the graph, we next needed a metric to evaluate whether T cells exhibit biased movement towards the parasite. However, as far as we are aware there have been no methods developed to detect bias in walks on graphs nor to simulate biased walks on graphs.

We previously developed rigorous methodology to detect attraction of cells moving in open space/3D towards a specific location [34]. Our test was based on comparing the distribution of angles between the movement vector and the vector to the infection site to the von Mises-Fisher (**VMF**) distribution (**eqn. (2)**), allowing us to detect deviations of the angle distribution from uniform [34]. We also used the VMF distribution to simulate biased random walks including correlated random walks [34, 49]. Now, we extended this work to develop a methodology to simulate biased random movement of T cells on graphs.

First, for every graph vertex, using a method based on Dijkstra’s algorithm we calculated the shortest path weight (**SPW**) that is the smallest sum of all edge weights from the vertex to any of the “infected” vertices [40]. A T cell that is attracted to the infection site should then on average reduce SPW values of vertices the cell visits over time (and eventually should reach one of the “infected” vertices). By analogy with the VMF distribution, we defined the probability for a T cell move between vertex *i* and *j* as *P_ij_* with a parameter *α* denoting the strength of attraction of T cells towards the infection site (**eqn. (4)** and **Figure 2**). For *α >* 0, cells have a higher probability of moving closer to the parasite (leading to decrease in SPW) than staying in place or moving farther (**Figure 2**). By calculating vertices visited by a T cell in a simulation, we propose a graph-based metric/test that allows to determine statistically if the cell movements deviate from unbiased (**eqns. (5)–(6)**).

One important feature of liver-localized CD8 T cells is their motility – these cells typically move with average speeds of 5 *−* 20 *µ*m*/*min [26, 27, 38, 53]. In our initial simulations of movement of unbiased T cells (with *α* = 0) we found that the cells moved between vertices less frequently than we observed in experiments; therefore, we introduced a motility parameter *ψ* that determines the likelihood of a cell for a given time step to stay at the same vertex, compared to moving to another vertex given the edge weight connecting the vertices (**eqn. (10)**); negative values of *ψ* decrease the probability of the cell to stay in the current vertex, thus, increasing the overall speed of a cell (**Supplemental Figure S4**). When simulating movements of cells from a particular experiment we chose values of *ψ* so we could approximately match the distribution of speeds and pauses observed in the data (e.g., **Supplemental Figure S5**). Simulated cells also displayed a high degree of persistence (as determined by a large proportion of small turning angles, **Supplemental Figure S5C**) that is consistent with other experimental data [38].

We next sought to investigate how well our novel graph-based test is able to detect biased movement of T cells in simulations. In these short (up to 3 h of movement time) simulations we used graphs derived from movie CTV5 (**Table 1**) along with movement characteristics of liver-localized CD8 T cells (**Supplemental Figure S5**) and the location of the “infected” vertices (**Supplemental Figure S1**). We randomly selected a vertex for the initial position of a T cell and then simulated its movement for a number of time steps (typically 300) assuming attraction strength *α* and motility *ψ* (**Figure 2** and see Materials and methods for detail). For every cell we then used information on vertices visited and used the likelihood ratio test to evaluate whether a T cell is detected as attracted towards the infection, both using open space/3D- and graph-based metrics (**eqn. (3)** and **eqn. (6)**, respectively). We repeated the simulations for 1,000 cells (**Figure 3**).

**Figure 3:**
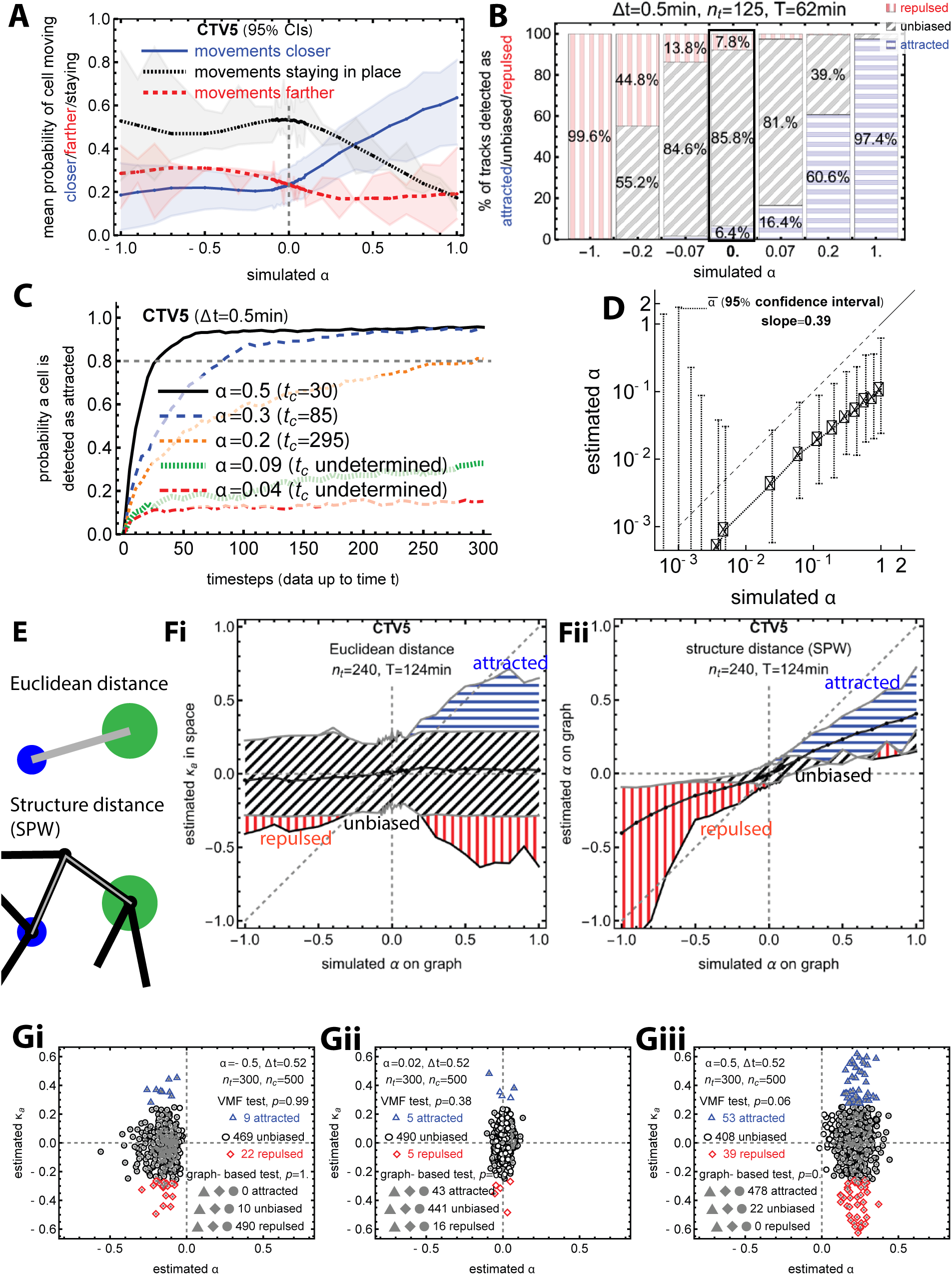
A new graph-based test for biased movement correctly detects moderately attracted cells as attracted and pseudorandomly moving cells as unbiased. We simulated movement of 1000 cells on a graph generated from liver sinusoids (experiment CTV5, see **Table 1**) for 50 values of *α* between −1 and 1 with a timestep Δ*t* = 0.5 min. **A**: Proportions of movements getting closer to (blue, SPW decreasing) or farther (red, SPW increasing) from the parasite or staying (black, SPW not changing) along with 95% confidence intervals (shaded area). **B**: Proportions of cells detected as attracted (blue, horizontal dashing), repulsed (red, vertical dashing), or unbiased (gray, diagonal dashing) by our new attraction test (**eqn. (6)**). **C**: Probability to detect a cell as attracted (*α >* 0) with our graph-based test as a function of the number of timesteps in a movie, with the critical *t_c_* defined as at least 80% of cells detected as attracted. **D**: Estimated attraction parameter *α* from **eqn. (6)** for different values of assumed parameter *α* is lower by about 60%. **E**: Schematic illustrating differences in the path assumed that a cell would take if moving in open space (Euclidean distance) or in a structure. **F**: For each simulation we calculate the T cells’ attraction towards the parasite assuming that cells move in open space (concentration parameter *κ_a_* of the VMF distribution, see **eqn. (2)** and **Fi**) or assuming that cells move via sinusoids (attraction parameter *α*, see **eqn. (6)** and **Fii**). We highlight cases when the estimated parameter is statistically different from zero (attracted: red, vertical dashing, repulsed: blue, horizontal dashing) or when cells are unbiased (black, horizontal dashing). **G**: We compared the relationship between the estimated concentration parameter *κ_a_* (open space) and the estimated attraction parameter *α* (structure) for cells moving in the structure with repulsion (*α* = *−*0.5, Gi), no bias (*α* = 0, Gii) or attraction (*α* = 0.5, Giii) towards the infection. Cells are styled based on the test results from the VMF test, detected as attracted (blue, triangles), repulsed (red, diamonds), or unbiased (black, circles). Cells detected as significantly biased by the graph-based test (attracted or repulsed) are filled gray.

Results of these “short” simulations showed that our proposed methodology of modeling biased random movements and detecting attraction on graphs works relatively well (**Figure 3**). In particular, the higher the values of the attraction parameter *α*, the more likely a cell is to move towards the parasite; however, because of the randomness of simulations even with a larger *α*, the cell can still move away (**Figure 3A**). In the case of no attraction, very few cells are detected as biased (attracted or repulsed) suggesting that our graph-based method (**eqns. (5)–(6)**) is unbiased (**Figure 3B**). Furthermore, for *α* = 1, the test accurately detects *>* 97% cells as attracted (**Figure 3B**). However, running simulations for different amounts of time revealed that there are limits on attraction strength *α* that can be detected by our graph-based metric; in particular, to detect an attraction strength of *α* = 0.2, nearly 300 time steps are required (**Figure 3C**). If a time step is 30 sec, detecting *α* = 0.2 for an individual cell would require over 2 hours of data collection. Finally, we detected a consistent bias in detecting attraction strength *α* given the simulation attraction strength *α* (**Figure 3D**, the detection is only 40% of assumed strength), further highlighting the challenge of accurately estimating *α*. This means that cells with weak attraction may be detected as unbiased, which may be unavoidable with statistical testing.

We previously showed that a metric based on a VMF distribution is statistically most powerful at detecting attraction of agents moving in open space/3D [34]. However, since liver-localized CD8 T cells do not move in open space and are constrained by the liver sinusoids, we next sought to test if a VMF-based metric (**eqn. (2)**) is able to accurately detect attracted cells when the cells move in a constrained environment, i.e., on the graph (**Figure 3E**). In our data we found that the path from a given T cell to the infection site is about 1.5 times longer via sinusoids than via direct Euclidean distance (**Supplemental Figure S6**) suggesting that methods to detect attraction in open space/3D may not perform well on realistically constrained movements. Interestingly, in simulations where we varied T cell attraction strength towards the infection site *α*, the open space/3D-based metric allowed us to detect cells as attracted only at very large values of *α*; in contrast, the graph-based test detected cells as attracted at relatively low simulated attraction strengths (**Figure 3F**). Furthermore, simulating movements of larger numbers of cells further demonstrated the lower power of the VMF-based test at detecting cells as attracted even with relatively high attraction strength *α* (**Figure 3G**). We hypothesize that the lower power of the VMF-based metric at detecting attraction of moving T cells on a graph arises because of the discretization of the space – if a T cell stays in the same vertex, this “no-movement” is uninformative for the open space/3D-based metric; however, for a graph-based test, staying in the same vertex may imply absence of attraction (**Figure 2**). Thus, our analysis strongly suggests that while the VMF-based metric may detect no attracted T cells in actual experiments, the graph-based metric should be more sensitive at detecting attraction if it exists.

While our novel graph-based metric (**eqn. (5)**) allowed us to detect attracted cells in simulations, it required a moderate level of attraction strength (*α ≥* 0.2) and a large number of recorded movements (*n_t_* = 300, **Figure 3B&C**). We therefore investigated if alternative graph-based metrics may have a higher power at detecting attracted cells. Interestingly, two metrics: monotonicity in SPW change (dubbed as “monotonicity”) with time or average change in SPW between initial value and all other values (dubbed as “T-test”) allowed us to detect cells as attracted at smaller values of the simulated attraction strength *α* (**Supplemental Figure S7A**). However, with a greater sensitivity, these two metrics had a high false-positive rate, detecting 50% of cells as attracted in simulations with unbiased cells (with *α* = 0, **Supplemental Figure S7A**). As expected, the VMF-based metric performed poorly and detected only a small proportion of cells as attracted (**Supplemental Figure S7A**). A “combo” test that compares the difference in the number of cells detected as attracted vs. repulsed (**eqn. (8)**) performed similarly to our main graph-based metric (**Supplemental Figure S7A**). Finally, we found that it is easier to detect attraction of a collection of cells all of which are assumed to be attracted to the infection site; we can robustly detect 100 cells as attracted if their assumed level of attraction strength is *α* = 0.2 (**Supplemental Figure S7B&C**). Thus, out of the tested metrics, our attraction-on-graph metric (**eqn. (5)**) is the most robust metric to detect attraction of moving T cells toward Plasmodium liver stages.

### A higher proportion of T cells is detected as attracted to the infection site in open space/3D but not via sinusoids

After having developed a rigorous methodology to detect biased movement of T cells on graphs, we analyzed our new data on the movement of liver-localized CD8 T cells in their search for Plasmodium liver stages (**Figure 1** and **Table 1**). Our three novel experiments followed T cell movements in distinct scenarios, i.e., when T cells were just approaching the parasite (CTV3), when we observed a cluster of few (2-3) T cells (CTV5), and when we observed a very large T cell cluster (10 cells, Mouse 4, **Tables 1 and 2**). As in our previous work, T cells in these movies exhibited rapid correlated random walks with most turning angles being small indicating directed movement (**Supplemental Figure S5Ai&Aiii** and [38]). Also consistent with our previous work [34], when using the open space/3D attraction metric (**eqn. (2)**), we found that most cells do not display bias relative to the infection site, and only in the presence of a moderately sized cluster (movie CTV5) did the number of T cells detected as attracted nearly statistically exceeded the number of cells detected as repulsed (**Table 2** and **Figure 4Ci&Di**). Similarly, with the same metric we found that there is a small bias of all moving T cells towards the infection site in the presence of a small T cell cluster, but not in the situation where no T cell found the parasite - and surprisingly, neither when there was a supercluster of T cells (**Supplemental Figure S8**). These results recapitulate our recent finding that only a small proportion of T cells searching for the liver stage display detectable (with VMF distribution-based metric) attraction towards the infection site [34].

**Figure 4:**
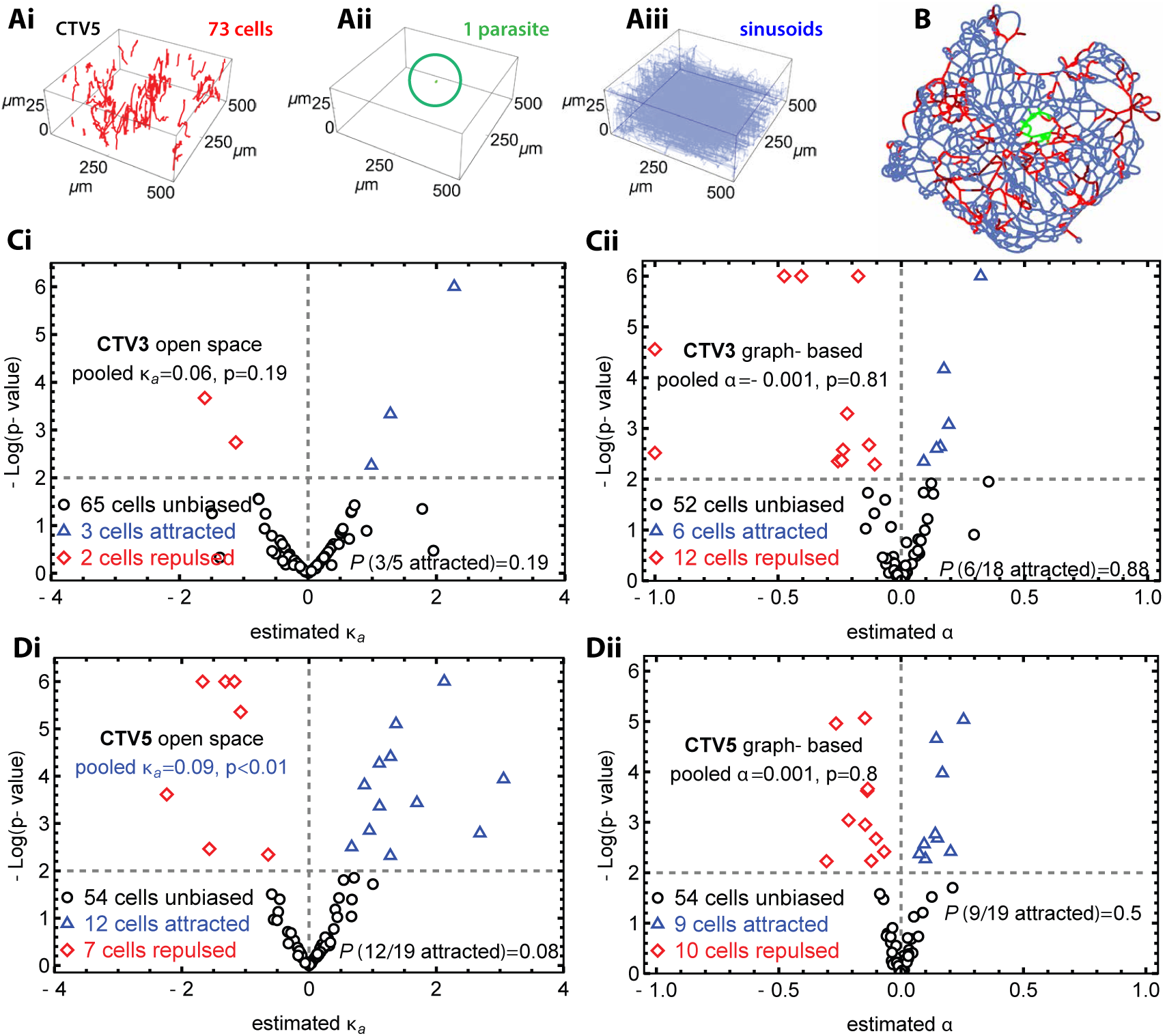
Plasmodium-specific CD8 T cells exhibit weak attraction towards the liver stage in open space/3D but not via liver sinusoids. We performed experiments in which we tracked the movement of liver-localized Plasmodium-specific CD8 T cells in liver sinusoids after infection with Plasmodium sporozoites (Figure 1) and quantified the attraction of T cells to the infection using either the VMF distribution (**eqn. (2)**, open space metric) or our graph-based test (**eqn. (6)**). **A**: We show trajectories of T cells (**Ai**), the position of the parasite/liver stage (**Aii**), and digitized liver sinusoids (**Aiii**) generated from one of the experimental movies (CTV5). Note that axes are not to scale due to the shallow nature of imaging experiments. **B**: By combining all elements in **A**, we represent sinusoids as a graph and map T cell and parasite positions to vertices on the graph (see Figure 1 and Materials and methods for more detail). **C-D**: For each T cell in the dataset CTV3 (no T cell cluster, see **Table 1**, **C**) or CTV5 (small T cell cluster, **D**), we calculated the concentration parameter *κ_a_* (**i**) and attraction parameter *α* (**ii**), measuring T cell attraction towards the infection site in open space and on the graph, respectively. The results are presented as volcano plots with the estimated parameters and corresponding p values from LRT. Results of analysis of the data from experiment Mouse4 were similar to above, and additional estimates of T cell bias relative to the infection site are listed in **Table 2**. Bias in the number of cells detected as attracted vs. all biased cells was evaluated using the binomial distribution (**eqn. (7)**).

**Table 2:** CD8 T cells are detected as attracted to the infection site when using the open space/3D metric but not when using the graph-based metric. For three of our datasets where CD8 T cells, sporozoites, and liver sinusoids have been imaged in one setting, we calculated the proportion of cells detected as attracted, repulsed or unbiased using either the open space/3D metric (based on the VMF distribution, **eqn. (2)**) or using our novel graph-based metric (**eqn. (6)**, see **Figure 4**). We also calculated the concentration parameter *κ_a_* from VMF distribution and the attraction parameter *α* for all cells (“pooled” estimates) and corresponding *p* values from the LRT (see Materials and methods for detail).

| Experiment<br>cluster size (# T cells/liver stage) | CTV3<br>0-1 | CTV5<br>2-3 | Mouse4<br>10 |
| --- | --- | --- | --- |
| #attracted in 3D (VMF) | 3 | 12 | 2 |
| #repulsed in 3D (VMF) | 2 | 7 | 3 |
| #unbiased in 3D (VMF) | 65 | 54 | 49 |
| binomial test $p$ | 0.19 | 0.08 | 0.55 |
| $\kappa_a$ for all cells | 0.059 | 0.087 | 0.04 |
| $p$ | 0.19 | <0.01 | 0.22 |
| #attracted via sinusoids | 6 | 9 | 3 |
| #repulsed via sinusoids | 12 | 10 | 1 |
| #unbiased via sinusoids | 52 | 54 | 50 |
| binomial test $p$ | 0.88 | 0.5 | 1.0 |
| $\alpha$ for all cells | -0.001 | 0.001 | -0.004 |
| $p$ | 0.81 | 0.80 | 0.5 |

Our simulation results strongly suggested that for T cells moving on graphs, our novel graph-based attraction metric (**eqn. (5)**) is more sensitive and robust at detecting attracted cells as compared to open space/3D (VMF)-based metric (**Figure 3F&G**). Therefore, to apply the graph-based attraction metric we converted the 3D coordinates of moving T cells to positions on liver sinusoids-derived graphs (**Figure 1**). Specifically, we carefully mapped 3D positions of every cell at every time point to the nearest vertex of the subdivided graph representing liver sinusoids (**Figure 4A&B**). Because the parasite is located inside of a hepatocyte, the infection site comprises of hundreds of vertices, denoted as “infected” vertices (**Figure 4B** and **Table 1**). As noted earlier, the actual distance between T cells and the infection site via liver sinusoids (or on the graph) is around 50% longer than the Euclidean distance (**Supplemental Figure S6**) suggesting further limitations of open space/3D-based metrics to evaluate biased movement of T cells.

We then collected vertices visited by every T cell during the imaging, calculated the probability that the T cell was getting closer to the “infected” vertices, and determined if the estimated attraction strength *α* is statistically significiantly different from 0 (**eqn. (6)** and see Materials and methods for detail). Surprisingly, we found similar proportions of T cells detected as biased (18 cells for CTV3 and 19 for CVT5) as with the VMF distribution-based metric (**Figure 4C&D** and **Table 2**). But in contrast to the results of the VMF-based metric, we found similar numbers of T cells detected as attracted or repulsed via sinusoids, suggesting that detecting cells as attracted may be false positives (**Table 2** and **Movie 5**). Pooled estimates of the attraction strength *α* for all T cell movements also revealed no evidence of attraction (**Figure 4Cii&Dii** and **Table 2**). Importantly, cells detected as attracted demonstrate a reduction in the SPWs of the vertices the cells visit and cells detected as repulsed demonstrate an increase in the SPW over time, suggesting that the graph-based metric correctly identifies biased cells (**Supplemental Figure S9**). However, because we found similar numbers of T cells detected as repulsed or attracted and a lack of a strong estimated attraction strength *α* for all cells (**Table 2**), detecting the moving T cells as biased is likely to arise as an artifact, and detection of attraction via the VMF-based metric may also be an artifact.

We wondered if finding similar numbers of attracted and repulsed cells and lack of overall bias in estimated attraction strength *α* could be a result of converting 3D images of liver sinusoids to graphs whereby we force nearby 3D positions to be mapped to the same vertex. This may have the effect of eliminating minuscule movements of cells (on average, less than 3 *µ*m) from consideration. The open space-based VMF distribution-based metric, by using actual 3D positions of T cells, has no such guard rails against ignoring negligible movements. To check this, we reran the VMF test using positions in space collapsed to a 3D grid with side length 3 *µ*m, which lowers the numbers of cells detected as attracted but does not eliminate preferences for attraction in clustered datasets (compare **Supplemental Figure S10i** to **ii**) in both our current data (**A**) and our previously published dataset (**B**, [34]). Thus, lack of attraction of liver-localized T cells towards the infection site is not due to converting images of liver sinusoids to graphs.

When comparing results of the analysis of experimental data with stochastic simulations of T cells moving on graphs we noted that we detected more cells as attracted in experiments — in particular, we found between 6 to 12% of moving T cells to be attracted to the infection via liver sinusoids but under 2% of T cells were detected as attracted in simulations (**Supplemental Figure S11**). This suggested that perhaps there are some elements of how T cells move in liver sinusoids that may be not fully represented in simulations. Alternatively, there may be something special about liver areas where the parasite is located, e.g., perhaps areas of higher traffic may increase the chances of T cells to be detected as attracted.

### Random/unbiased search of T cells for the infection site via liver sinusoids can be surprisingly efficient

Our finding that liver-localized CD8 T cells searching for Plasmodium liver stages did not display a significant bias in movement towards the infection site was surprising, especially given previous theoretical arguments that too many unbiased T cells would be needed to survey the whole liver and eliminate all liver stages in 48 h [26, 34]. However, these previous theoretical models assumed that CD8 T cells search for infection in open space and/or a regular lattice; previous theoretical results suggested that search in constrained environments may be quite efficient [38, 39]. Because it takes 44-48 hours for the liver stage in mice to mature [8, 54], T cells moving in liver sinusoids may in fact have sufficient time to find the infection. Therefore, we next simulated T cell search for the infection on the longer time scale (up to 48 hours of infection).

To run stochastic simulations determining the time it would take for a T cell to find an infection site on liver sinusoids-based graphs, we first needed to determine how many T cells would typically be detected in the imaging volume. Livers of adult mice have a volume of 0.8 cm^3^ = 0.8 *×* 10^12^ *µ*m^3^ [55]. The imaging volume is about 512 *×* 512 *×* 46 = 1.2 *×* 10^7^ *µ*m^3^, which is about 1*/*66, 343 = 1.5 *×* 10*^−^*^5^ of the entire liver. Based on a review of previous studies, a minimum of 1 *−* 2 *×* 10^4^ cells in the liver are sufficient to protect the mice from a challenge with Plasmodium sporozoites [32], corresponding on average to 0.2-0.3 liver-localized CD8 T cells per imaging volume. In our experiments, around 10^4^ *−* 10^5^ T cells accumulate in the liver after adoptive transfer of activated CD8 T cells [43] which may be on the higher end of protective T cell numbers. In our long-term simulations we therefore opted to have on average 1 CD8 T cell per imaging volume, corresponding approximately to 6.25 *×* 10^4^ liver-localized T cells.

One issue with simulating T cell movement on a graph is that eventually the cell will reach a boundary vertex located at the edge of the imaging volume, and with the next movement, may then leave the imaging volume. When the cell reaches a boundary vertex and remains at boundary vertices for two time points in a row, we consider it to have left the imaging volume; a new cell then enters the imaging volume starting on a pseudorandomly selected boundary vertex. Therefore in a simulation of 1 cell search for the parasite, there is exactly 1 cell in the imaging volume at each time, but in 48 hours as many as 1000 “cells” leave and re-appear in the imaging volume. Most re-appearing cells in fact leave the imaging volume again, because a cell at a boundary often stays at the boundary for several time steps and is thus modeled as immediately leaving.

Having set all prerequisites, we next ran simulations of a single T cell search for an infection site by varying the attraction strength *α* from −1 (strongly repulsed from the infection site) to 1 (strongly attracted to the infection site) for 48 hours (using the graph and T cell movement length/displacement distribution from movie CTV5). We stopped individual runs if a T cell moved to an “infected” vertex or if the time reaches a 48 h threshold (**Figure 5**). As expected, the attraction strength had a major impact on the time and probability that a T cell finds an infection site, e.g., for *α >* 0, the T cell always found the infection site within 48 hours and for larger *α* it took only a few hours for the T cell to find an infected vertex (**Figure 5A**). However, unexpectedly, we found that in the absence of attraction (*α* = 0) or even when T cells were repulsed from the infection site (*α <* 0) a T cell was successful at finding the parasite with *>* 94% probability, and within 10-12 h of search time (**Figure 5A**). Additional power analysis suggests that a single unbiased (*α* = 0) T cell (per imaging volume) has about 80% chance of finding the infection site in 24 h (**Figure 5B**). We found similar results for graphs and cell displacement distributions from movies CTV3 and Mouse 4 (not shown).

**Figure 5:**
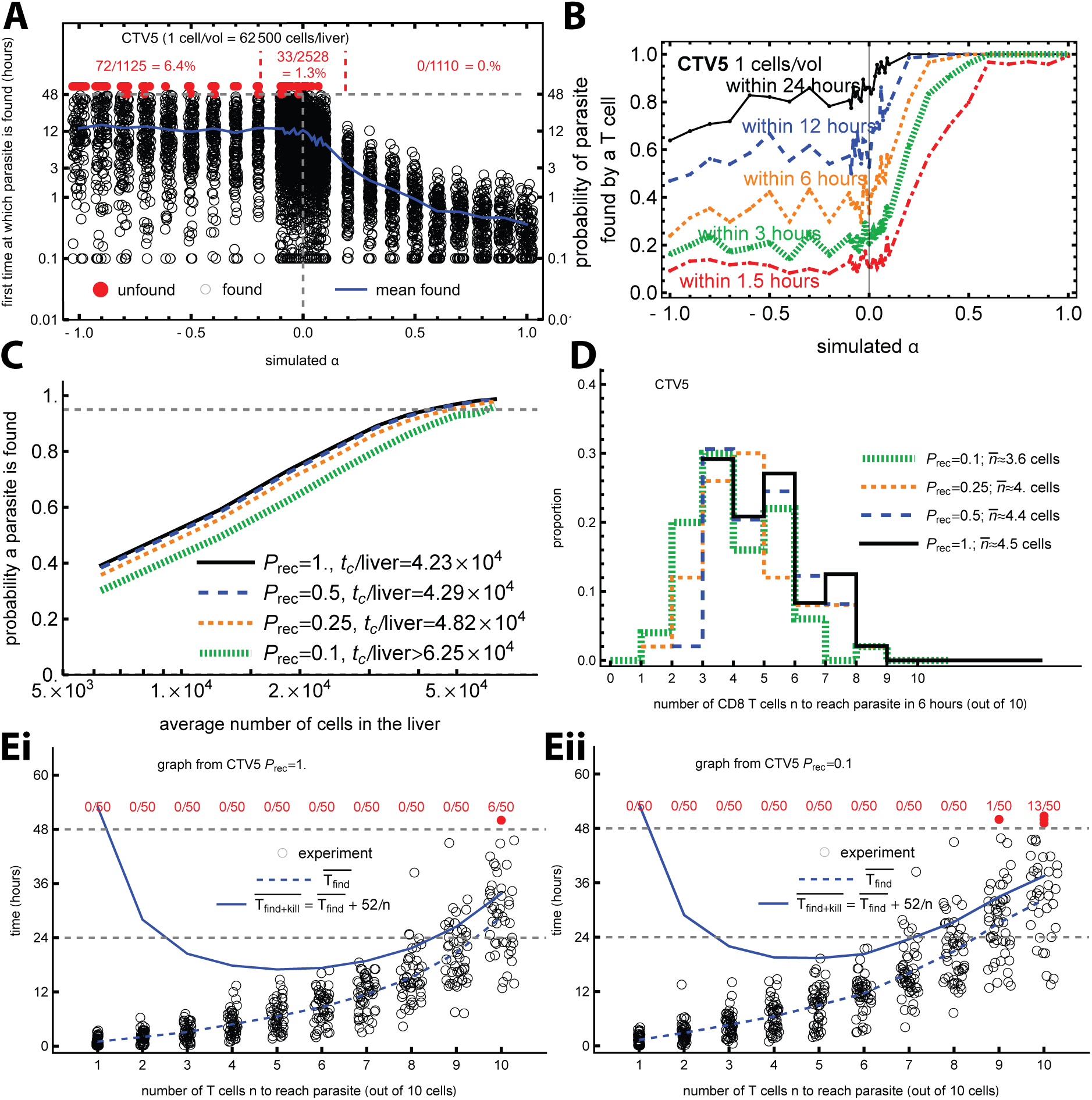
A relatively small number of liver-localized unbiased CD8 T cells is sufficient to locate a liver stage within 12 hours. We simulated movement of T cells on the graph (based on experiment CTV5) searching for an infection while varying the attraction coefficient *α* and calculating the time until T cells find the infection. In simulations we assumed 1 T cell per imaging volume, corresponding approximately to 6.25 *×* 10^4^ T cells in the liver (see main text for more detail) with speeds and persistence of the movement matching experimental data, with Δ*t* = 0.5 min (**Supplemental Figure S5**). **A**: Time to find the infection for different values of *α* out of 100 simulations. Red points indicate runs for which the parasite was not found within 48 hours. The blue line indicates the average time at which the parasite is found for different values of *α*. **B**: Probability that T cells find the parasite within different times after infection as a function of the attraction parameter *α*. **C**: Probability that T cells will find the parasite as a function of the number of liver-localized CD8 T cells, with the critical number of cells per liver *t_c_* defined as the minimum number of cells for which 95% of parasites are found. We also introduced a probability *P*_rec_ that a T cell recognizes the parasite upon reaching an “infected” vertex. **D**: Number of T cells that will find the parasite after 6 hours. We simulated movements of 10 T cells per imaging volume (50 simulations per parameter set) with *α* = 0 (no attraction) for 6 hours and assumed that if a T cell recognizes the parasite, it stops moving. We also varied the probability that a T cell recognizes the infection *P*_rec_. **E**: For the same simulations from **D**, we calculated the time taken for different numbers of T cells (out of 10) find the parasite (T_find_) assuming *P*_rec_ = 1 in **Ei** or *P*_rec_ = 0.1 in **Eii**. Given our recent estimate that *n* T cells kill the para<u>site in</u> *<u>T</u>*<u>_k_</u>_ill_ = 52*/n* h [33], we also plot the average total time it would take for 10 cells to find and kill the parasite T_find+kill_ (blue line). Results were similar when assuming 0.3 cells/volume (or 2.3 *×* 10^4^ cell/liver, **Supplemental Figure S12**).

It is interesting that for “repulsed” T cells (with *α <* 0), the average time to find the parasite was apparently independent of *α* (**Figure 5A**). This may be a consequence of implementing one T cell per imaging volume because i) cells fleeing the parasite will invariably reach boundaries and then sometimes pseudorandomly may re-appear at another boundary vertex that appears to be closer to the parasite, and ii) the average time to find the infection site is taken only from cells that do arrive, so any cell that takes longer to find the infection site is excluded from calculating the average.

Our calculation suggests that one T cell per imaging volume correspond to about 6.25 *×* 10^4^ cells in the whole liver. Some previous studies found that an even smaller number of liver-localized CD8 T cells may be protective [56, 57]; to explore having on average fewer than one T cell per imaging volume, we artificially added empty time after a cell leaves the imaging volume. For example, if we want 0.5 cells per imaging volume (corresponding to 31,250 cells in the entire liver), then every time a cell leaves the imaging volume, we wait the same amount of time the cell moved before introducing a new cell at another pseudorandomly chosen boundary vertex. Running long simulations, we found that even when there are only 0.3 cells in the imaging volume, unbiased T cells (with *α* = 0) have a 73% chance of finding the infection site within 48 h, and power analysis suggested that unbiased T cells have 50% chance of finding the infection in 24 h (**Supplemental Figure S12**).

Given high and yet variable efficiency of T cell search for a liver stage on liver sinusoids-derived graphs (**Figure 5A**), we next explored if changing maximal speed of T cells or the number of parasites T cells need to searching for may impact the overall ability of T cells to locate all infected hepatocytes in the liver. We therefore performed additional simulations by restricting the maximum speeds T cells may exhibit; in part, this is because some of the rapid displacements that we detected could be floating with the blood flow events [27, 38, **Supplemental Figure S5Ai**]. Importantly, restricting maximal speeds to be less than 40 *µ*m*/*min had minimal impact on the average time (or its variance) to find the infection (**Supplemental Figure S13A**). In contrast, the average time needed for a single T cell/volume to find parasites located in different parts of the liver increased with the number of parasites, and for 20 or more parasites/liver, randomly moving T cells had only 60% chance to locate all parasites in 48 hours (**Supplemental Figure S13B**). This suggests that more than 6 *×* 10^4^*/*liver would be required for randomly moving T cells to find multiple (*>* 20) parasites in murine liver.

So far, our simulations focused on the scenario of T cells finding the infection site (i.e., one of the infected vertices, **Table 1**). However, in order for a T cell to eliminate a parasite, the T cell must recognize that the hepatocyte is infected. This might not occur when the T cell first reaches an infected vertex. We therefore ran another set of simulations in which we introduced a probability of recognition *P*_rec_ per time frame that a T cell, upon reaching an infected vertex, recognizes that the hepatocyte is infected. In these simulations we varied the number of T cells per imaging volume (with scaling such that one cell per imaging volume is 62,500 cells per liver) and calculated the probability that a T cell finds and recognizes the infection in 48 h. Interestingly, about 4.23 *×* 10^4^ of liver-localized CD8 T cells are needed to find the infection in 48 h with 95% probability when *P_rec_* = 1, and this number increases only moderately to 4.82 *×* 10^4^ per liver when *P_rec_* = 0.25 (**Figure 5C**). The weak dependence of the number of T cells needed to find the infection site on *P_rec_* arises in part because when a T cell reaches the infection site, it will not leave it for a while due to many vertices being “infected”, so even if the infection is not recognized immediately, it will be recognized in a few time frames.

Given the relatively high efficiency at which unbiased T cells find the infection site, we wondered: if there are multiple T cells per imaging volume, would they be able to find the parasite early and form clusters? Clusters around a Plasmodium liver stage consisting of 3-6 CD8 T cells, and sometimes as many as 10-15 CD8 T cells, have been observed in experiments and explained best with the DDR model [31, 32, 58]; in this model, the first T cell searches for the infection site randomly but after finding the infection site, recruits other T cells. We therefore ran simulations assuming 10 unbiased (*α* = 0) CD8 T cells per imaging volume (with a cell that reaches a boundary vertex and leaves reappearing at a randomly chosen boundary vertex) searching for infection site for 6 h. By assuming that if a T cell recognizes the infection it stops moving, we found that out of 10 searching T cells, between 1 to 9 cells would find the infection site in 6 h (**Figure 5D**). This result suggests that even unbiased T cells have a remarkable search efficiency in liver sinusoids-derived graphs and challenges our previously established DDR model as the best model to explain T cell clustering data [32].

Finding the infection site and recognizing that a hepatocyte is infected are important, but they are not the only steps required for CD8 T cells to provide protective immunity — T cells must also kill the parasite before liver stages release blood-stage merozoites into circulation. With intravital microscopy, we previously tracked how clustered CD8 T cells eliminate Plasmodium liver stages, and with mathematical models, we quantified the killing efficacy of Plasmodium-specific CD8 T cells [31, 33]. We found that killing of liver stages by T cells follows the law of mass-action, where the rate of parasite’s death is proportional to the number of clustered CD8 T cells *n*, and that the time required to eliminate the LS is *T_kill_ ≈* 52*/n* h [33]. We ran long (48h) simulations assuming that there are 10 unbiased (*α* = 0) CD8 T cells per imaging volume (using parameters from movie CTV5) and calculated the time it takes for different numbers of T cells to find, recognize, and kill the parasite (*T*_find+kill_) with the probability of a T cell recognizing the vertex as “infected” *P_rec_* being either high (*P_rec_* = 1) or low (*P_rec_* = 0.1, **Figure 5E**). As expected, the time needed for *n* T cells to locate the infection site increased with *n*, and more than 24 h on average is needed for all 10 T cells to find the infection site; the time was again moderately dependent on the ability of T cells to recognize the infection *P_rec_* (**Figure 5**Ei&Eii). However, because the time required to kill the parasite declines with the number of clustered T cells, the overall time to find and kill the parasite is minimized for clusters of intermediate size, 3-8 T cells (**Figure 5E**). Thus, our simulations suggest that liver-localized CD8 T cells when present in sufficient numbers (*∼* 10^5^/liver) are capable of finding and killing all liver stages without a need for an attraction towards the infection site.

## 4 Discussion

CD8 T cells can be very efficient at eliminating intracellular infections, and yet, how these cells find and eliminate pathogen-infected cells remains poorly understood. It is generally believed that random, unbiased search for rare targets may be inefficient [26, 34, 59] and some studies suggest that attraction of T cells to sites of infection (or inflammation), driven by chemokines, is important for the efficiency of T cell responses [60, 61]. Here we extended our previous work and investigated how liver-localized CD8 T cells search for Plasmodium sporozoite-infected hepatocytes in murine livers while considering the constrained structure of the liver sinusoids. With intravital microscopy, we carefully tracked the location of the parasite and CD8 T cells over time, along with the structure of liver sinusoids in which liver-localized CD8 T cells move (**Figure 1**). We developed a novel pipeline to convert 3D images of liver sinusoids into weighted graphs and to assign 3D positions of the parasite and T cells to vertices on these graphs (**Figure 1**). We developed novel methodology to simulate the movement of T cells on these graphs with varying levels of attraction (or repulsion) towards the infection site (**Figure 2**). We also developed a novel metric to detect attraction of T cells moving on weighted graphs towards a collection of infected vertices and showed that this graph-based metric can accurately detect attracted cells (**Figure 3**). Surprisingly, with this new graph-based metric we found similar proportions of liver-localized CD8 T cells detected as attracted or repulsed (yielding no evidence of bias in movement for all cells) even though T cells appeared to be attracted to the infection site (when using open space/3D-based metric (**Figure 4** and **Table 2**) with few T cells clustered around the parasite. Stochastic simulations of unbiased T cells, moving on liver sinusoids-derived weighted graphs, suggested that even unbiased T cells (with attraction strength *α* = 0) are able to locate and eliminate the parasite within 24 h of infection (**Figure 5** and **Supplemental Figure S12**). Thus, while we do not find evidence that liver-localized CD8 T cells are attracted to the infection site via sinusoids, simulations suggest that such attraction is not required for T cells to effectively locate and eliminate the parasite.

Many details of leukocyte movement in 3D and in constrained environments are well understood but some mechanisms, e.g., how cells move their nuclei via small channels remain debated [62–67]. Previous studies addressed the movement of cells via artificially designed channels that may influence the overall motility and speeds of cells [62, 63]; however, less is known about the impact of specific tissue constraints on cell movement *in vivo* and on the efficacy at which T cells locate the site of infection. Previous studies also modeled the blood vasculature of tissues [68, 69]; however, as far as we know our study is the first to convert digital images of the blood vessels into weighted graphs allowing us to apply a variety of graph-based algorithms to the analysis of such data and to the modeling of cell movement on the graphs [70].

Our findings that unbiased movement of T cells on the graphs with speeds and pauses observed experimentally (**Supplemental Figure S5**) is sufficient to locate the infection site within 48 h (**Figure 5**) may not be that surprising given that sinusoids constitute only 10-20% of the volume of the liver (**Supplemental Figure S2**), and thus, in sinusoids, T cells need to explore less space (as compared to the liver volume) to locate the infection. Furthermore, search for infection via liver sinusoids is mathematically not a search in open space/3D. It has been previously established that in a 3D lattice the probability of randomly finding a destination given infinite time is about 34%, while in 2D and 1D it is 100% [71]. We have used a notion of “fractional dimension” and calculated it for our liver sinusoid-derived graphs (**eqn. (1)** and **Table 1**). The fractional dimension was slightly above 2 for the original graphs and less than 2 for the subdivided graphs (**Table 1**). Therefore, a relatively high search efficiency of unbiased liver-localized T cells may simply arise due to a lower dimension space they have to explore.

Our work has several limitations. A typical issue with intravital imaging is the small volume of the tissue imaged and its closeness to the tissue boundary (typically within 40 *−* 50 *µ*m). We do not know if CD8 T cells behave similarly in deeper regions of the liver. We should note, however, that it is somewhat surprising that in our movies we typically observe 30-50 T cells per movie while calculations suggest that we should see *<* 10 cells per imaging volume. Whether there are more T cells near the liver boundary than in the deeper tissue remains to be determined.

We adoptively transferred in vitro activated CD8 T cells to generate liver-localized CD8 T cells prior to SPZ infection (**Figure 1A**). While this is not a vaccination aimed to induce liver-resident T cells (Trms), our previously studies found that in vitro activated CD8 T cells in the liver, imaged 2-6h after adoptive transfer, exhibit similar movement patterns as T cells, induced by vaccination with radiation-attenuated SPZs, and imaged 4 weeks post vaccination [27, 38]. Therefore, our finding that liver-localized CD8 T cells do not display detectable attraction towards parasites in the liver is unlikely to be due to the method of inducing T cells in the liver. However, future studies will need to investigate if vaccine-induced liver Trms also search for infection without attraction via sinusoids.

We used standard, well-developed tools in Imaris to process the imaging data for positions of the parasite and CD8 T cells, as well as to trace liver sinusoids (see Materials and methods for detail). However, automated tools in Imaris do produce false positives and require manual corrections. We typically used at least two different trained human operators to process and realign the objects in Imaris. We also took great care to correct for tissue drift by using the reference frame tool in Imaris. Yet, it is still possible that the results may be somewhat different if the original microscopy data was independently processed with a different software. Our imaging data did not allow us to rigorously evaluate the direction of blood flow in individual sinusoids. Blood flow may influence the ability of T cells to move via sinusoids even though previous studies typically did not find the blood flow direction to be important for liver-localized CD8 T cell movement [27].

We generated liver sinusoids-based graphs by using the length of sinusoids between two vertices as an edge weight, and assumed that T cells searching for infection site would take the path with the smallest SPW. However, other possibilities exist. For example, edge weights could be represented by the volume of the sinusoid between intersections. Also, the calculation of SPW does not take into account the number of different paths between the T cell and the parasite that a T cell could take that would have identical SPWs, and if there are many of such paths, moving along any of the paths would increase the chances of finding the parasite even if the specific movement does not decrease SPW. Methodologies to address these limitations will need to be developed. Given that our graph-based metric is more sensitive at detecting attraction of T cells on graphs as compared to an open space/3D-based metric (**Figure 3F&G**), we found it surprising that we did not find evidence of attraction of T cells in experiments with the graph-based metric but did detect attracted cells with open space/3D-based metric (**Figure 4**). We do not have a good explanation of why this may be the case. There may be a subconscious bias of the operator to choose a particular liver area for imaging, e.g., areas with subtle biased movements towards the infection that are negated by the sinusoids-based metric. Our previous work highlighted that the bias towards detecting more attracted cells increases with reducing the imaging volume [34]; however, this does not explain why we detect many attracted cells with the open space/3D-based metric in some movies (e.g. with moderate T cell cluster size) but not in others (e.g. with no clusters or very large T cell clusters). Ultimately, however, this suggests that detecting attraction of T cells towards an infection site with open space/3D metrics for T cells actually moving in an constrained environment may be misleading. Our previous studies suggested that following adoptive transfer of in vitro activated CD8 T cells, about 1% accumulate in the liver [43], and comparing the liver volume with the imaging volume suggested that on average we should typically see 1-2 T cells in the imaging volume. However, in our experiments we see at least 10 times more cells (**Table 1**). Whether the difference arises because there are more cells in the liver just under its surface, because when choosing to image a specific area one tries to focus on places with many T cells, or because the same cells may be leaving and re-entering the imaging volume remains to be determined.

An intrinsic assumption in our simulations is that all simulated cells have identical movement parameters (e.g., *ψ* and *α*); however, it is possible that some cells move with bias and other move randomly, as our previous work suggested [34]. Additionally, our simulated cells moving with attraction move with a constant attraction strength; if real cells do display attraction to the parasite, it likely would be dependent on the distance of the cell to the parasite. However, neither of these assumptions affect our conclusions about the time for cells to randomly find and kill the parasite. Another limitation comes in the form of an unexplained result: the numbers of cells detected as biased (i.e., attracted or repulsed) by the graph-based test is around 10 times more than expected, defining expectation as the proportion of cells detected attracted if cells all have the same *α* as the pooled *α* calculated from the dataset (**Supplemental Figure S11**). This could be explained if some cells truly moved with attraction and an equivalent number of cells move with repulsion, but we have assumed that cells do not move with repulsion in simulations of unbiased cells.

In our simulations we used the displacement of actual cells to simulate how T cells would move on graphs. Some of these displacements were relatively long (e.g., 50 to 100 *µ*m in 30 sec) and most likely represent floating events whereby T cells detach from the sinusoidal wall and float with the blood flow. We found that restricting speeds of moving T cells has negligible impact on the average time to find the parasite (**Supplemental Figure S13A**). Yet, simulations that use user-defined distribution of T cell displacements could be useful to further test how short and long displacements may influence efficiency of T cell search for the infection site in the liver. We assigned vertices that were within 40 *µ*m of the parasite’s location as “infected”, and reaching one of such “infected” vertices by a T cell was considered a successful search for the infection site. However, the exact size of the infected hepatocyte is not fully known. How the time required to find an infection depends on the chosen size of the infected hepatocyte remains to be investigated.

Finally, we only analyzed three novel movies in which we tracked the parasite, T cells, and labeled liver sinusoids in one setting. Therefore, the lack of T cell attraction towards the infection site in the liver via sinusoids may need to be tested in additional experiments. It is also unfortunate that our graph-based metric does not allow us to accurately estimate the simulated attraction strength; given that we only recover 40% of the simulated strength (**Figure 3D**) suggests that the lack of detecting most T cells as attracted in experiments could be due to a lack of power to detect weak attraction. Yet, as sinusoid structures are similar between experiments (**Supplemental Table S1**), our simulation conclusions that unbiased movement is sufficient to find and kill parasites within 48 h are not expected to change with more data.

Our work opens avenues for future research. Intravital imaging (with spinning disk confocal or two-photon microscope) typically allows to image a relatively small proportion of the liver (e.g., 0.001%) and extrapolating our results on how liver-localized CD8 T cells survey the whole liver would require data on the whole tissue. Previous studies have used confocal microscopy to visualize detailed structures of murine livers [72–75]; methodologies to rigorously quantify liver sinusoids (and other liver blood vessels) and convert them into graphs from such larger image sets will need to be developed. Of note, we expect that the whole liver would be represented by a subdivided graph with around 10^8^ vertices, which is an order of magnitude higher than a graph for the road network of the contiguous United States [76]. To more accurately map T cell positions to vertices on liver sinusoids-derived graphs. it is important to record the cell positions frequently. However, there is a trade-off between how frequently the imaging volume can be scanned and the overall duration of imaging; longer exposure to the laser over time may damage the tissue and render the generated data less useful. Using novel imaging techniques such as three photon microscopy may help generate movies with greater depth, and more frequent and longer imaging [77, 78]. Our conclusion that liver-localized CD8 T cells can effectively find for the infection site randomly, without strong attraction, suggests that the speed at which T cells move in the liver becomes an important factor in determining how many CD8 T cells are needed to eliminate the infection within the 48h of the liver stage lifespan. Yet, factors regulating T cell movement speeds in tissues remain poorly understood. Future studies should investigate why some T cells move faster than others, and if increasing the average speed of T cell movement in tissues may compromise their efficacy at scanning the tissues for infection. While we found that T cells may be very efficient at locating a single parasite in the liver, finding 10-20 parasites may take much longer time (**Supplemental Figure S13B**). Future studies will need to further quantify limits of CD8 T cell-based vaccines in settings of high malaria transmission when individuals receive many bites by infectious mosquitoes. Our results also suggest that analysis of cell tracking data requires better understanding of the tissue environment that is providing (or impeding) cellular movement. Detecting biased movement without taking into account underlying tissue structure may be a false positive result. Simulating T cell movement without accounting for specifics of tissue structure may also lead to erroneous conclusions. We hope that our novel framework to convert tissue images into weighted graphs and to simulate movement of biased cells on such graphs will be applicable to other cell and tissue types, and may explain why immune cells are able to control some and fail to control other infections.

## Data sources

All 3D coordinate data are available as a supplement to this paper and on github: https://github.com/viktorZenkov/sinusoids. Example Imaris files are available on zenodo: https://zenodo.org/records/5715658 and can be viewed in the free Imaris viewer.

## Code sources

Analyses have primarily been performed in Mathematica (version 13); several Mathematica-based scripts of how .hoc files are processed and how to run simulations of T cells on graphs are provided on github: https://github.com/viktorZenkov/sinusoids.

## Ethics statement

All animal procedures were approved by the Animal Experimentation Ethics Committee of the Australian National University (Protocol numbers: A2016/17; 2019/36). All research involving animals was conducted in accordance with the National Health and Medical Research Council’s Australian Code for the Care and Use of Animals for Scientific Purposes and the Australian Capital Territory Animal Welfare Act 1992.

## Author contributions

VSZ is the primary author of the paper, with broad supervision by VVG. VSZ developed methodologies for this study in collaboration with VVG. IAC performed imaging experiments. MAL oversaw the applications of graphs. RL processed imaging data in Imaris to trace cells and sinusoids, with additional processing by WNG and VSZ. VSZ performed analyses of the data, simulations of cell movements, and the dimension analysis. VSZ wrote the first draft of the paper that was then edited by VVG, with additional contributions by other co-authors.

## Acknowledgments

For long simulations, the following non-authors ran VSZ’s simulation code with parameters specified by VSZ on UTK’s HPCC (ISAAC) to create data more quickly: Saikat Bayabyal, Dipanjan Chakraborty, Kailynn Deck, John Maddox, Shivam Patel. We also would like to thank Estefania Hurtado for generating movies of how the original imaging data were processed in Imaris.

## Supplemental Information

### Additional experimental details

Mice were prepared for two-photon microscopy as previously described [27]. Mice were anesthetized with a mix of Ketamine (100 mg/kg) and Xylazine (10 mg/kg). Throughout the surgery and imaging procedure the mouse temperature was maintained at 37°C using a heating mat attached to a feedback probe inserted in the mouse rectum. A lateral incision was made over the left lobe of the liver and any vessels cauterized by applying light pressure to the vessel until clotting occurred naturally. The mouse was then placed in a custom made holder. The liver was then exposed and directly adhered to a cover slip that was secured in the holder. Once stable the preparation was transferred to a Fluoview FVMPE-RS multiphoton microscope system (Olympus) equipped with a XLPLN25XWMP2 objective (25x; NA1.05; water immersion; 2mm working distance). For analysis of the motility of the sporozoites and liver-localized CD8 T cells an approximately 50 *µ*m z-stack (2 *µ*m/slice) was typically acquired using a standard galvano-scanner at a frame rate between 1 to 2 frames per minute depending on the movie.

The selection of parasites was contingent upon clear, healthy sections of liver tissue, and the proximity of lymphocytes at the time of imaging. Each parasite chosen for imaging required unimpeded blood flow of surrounding sinusoids, bright autofluorescence and clear surrounding structures. In addition, the parasites chosen for imaging were also dependent upon what information we wanted. For the 0 hour timepoint, parasites were selected if they had healthy tissue and had no lymphocytes within a 40 *µ*m diameter of the parasite. This allowed for analysis of attraction of lymphocytes with no primary lymphocyte signaling from an infected cell. At 2 hours post infection, parasites were selected if they had healthy tissue and had *≥* 1 lymphocytes within a 40 *µ*m diameter of the parasite.

This allowed for analysis of attraction of lymphocytes with at least 1 primary lymphocyte signaling from an infected hepatocyte. We used the 80 *µ*m diamenter (*r* = 40 *µ*m) because it represents the average width of a standard murine hepatocyte [32].

### Imaging data processing details

Raw imaging data was analyzed using the Imaris x64 software (Bitplane), versions 9.8.2 to 10.0.1. Tracking of individual cells in a z-stack was performed using the “Spots” function in surpass mode. Extensive manual adjustments were subsequently done to ensure the accuracy of the tracks. The detection of individual cells relied upon their relative fluorescence intensity and size (diameter *≥* 9 *µ*m), and used the remaining default function settings for the calculation of the motion tracks using an autoregressive motion algorithm. Detection of fluorescence for each spot was manually adjusted to reduce background detection at each time point to ensure clear distinction of cell vs autofluorescence by the algorithm.

### Data formatting

To generate graphs from filaments produced by Imaris we use a .hoc (“HOC”) file as opposed to the usual statistics output of Imaris. This .hoc file contains a list of filaments, each of which includes points along the filament every 1 *µ*m, frequently enough to ensure that any bends are accounted for. It also contains information about every pair of filaments sharing a beginning or ending point. This information is sufficient for our later methodology that transforms a filament tracing into a graph. Exporting the .hoc from Imaris was done in the following way

- Go to File*→*Preferences or Edit*→*Preferences.
- Go to the Statistics tab.
- Ensure the “Filaments” checkbox has a full checkbox (not just a dash) and click OK.
- With the Filament object selected at the desired time to export a tracing, go to the Edit tab (labeled with an orange pencil icon).
- Click Export. One of the format options is “.hoc”. Choose that format and export.

### Graphs overview

A *graph* is a data structure which contains a finite number of *vertices* (also called nodes, represented as points) and *edges* (represented as lines). Edges connect pairs of vertices and may be directed (i.e. possessing a direction from one vertex to another) or undirected - we use undirected graphs in this work. Each edge may also have a numerical *edge weight*, allowing for weighted and unweighted graphs. Two vertices are called *adjacent* if there an edge connecting them in the graph.

A *walk* on a graph is a sequence of edges which join a sequence of adjacent edges. A *path* on a graph is a walk whose edges do not feature any duplicate vertices. The *shortest path* between any two vertices is the path which has the shortest length, where the length is the number of edges in the path in an unweighted graph and the sum of the edge weights of the path’s edges in a weighted graph. The *shortest path weight* (**SPW**) is then the accumulated weight of that path.

We note that sometimes, planarity is of interest in graph theory, i.e. the potential property that a graph may be able to be displayed in 2D without any edges crossing each other. Planarity is *not* of interest in this work, and the sinusoid graphs are not typically planar (even if they appear planar at first glance).

### Comparing graphs

When generating graphs from liver sinusoids in five independent experiments (and thus, different mice, **Supplemental Figure S1**), we wanted to establish that these graphs are in some way similar. Most graph comparison methods (like isomorphisms) are focused on matching individual unique vertices which is not fully applicable to our graphs because we are interested in comparing the general structure. Thus we used a distribution of *local clustering coefficient* (**lcc**), which is a measure of local density [79]. For each vertex *v*, the local clustering coefficient is

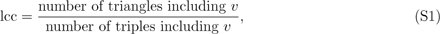

where a triangle is a trio of vertices which are all connected and a triple is a trio of vertices with at least two edges, both featuring *v*. Tallying the distribution of local clustering coefficients for each graph from our five movies gives five lists, which we compared pairwise using the Kolmogorov-Smirnov test (**Table S1**). All of the graphs are similar according to this test as all *p* values are nearly 1.

**Supplemental Table S1:** All liver sinusoids-derived graphs are statistically similar. For each of our five original (undivided) graphs (**Supplemental Figure S1**) we calculated distributions of local clustering coefficients (**eqn. (S1)**) and compared these distributions (pair-wise) using a Kolmogorov-Smirnov test. We show p values from that test.

| Experiment | Mouse C | Mouse D | CTV3 | CTV5 | Mouse4 |
| --- | --- | --- | --- | --- | --- |
| Mouse C | 1.0 | 1.0 | 1.0 | 1.0 | 0.999 |
| Mouse D | 1.0 | 1.0 | 1.0 | 1.0 | 0.995 |
| CTV3 | 1.0 | 1.0 | 1.0 | 1.0 | 0.999 |
| CTV5 | 1.0 | 1.0 | 1.0 | 1.0 | 0.998 |
| Mouse4 | 0.999 | 0.995 | 0.999 | 0.998 | 1.0 |

### SPW algorithm

To collect the set of SPWs from every vertex in the graph to the parasite, we used an algorithm inspired by Dijkstra’s algorithm [40]:

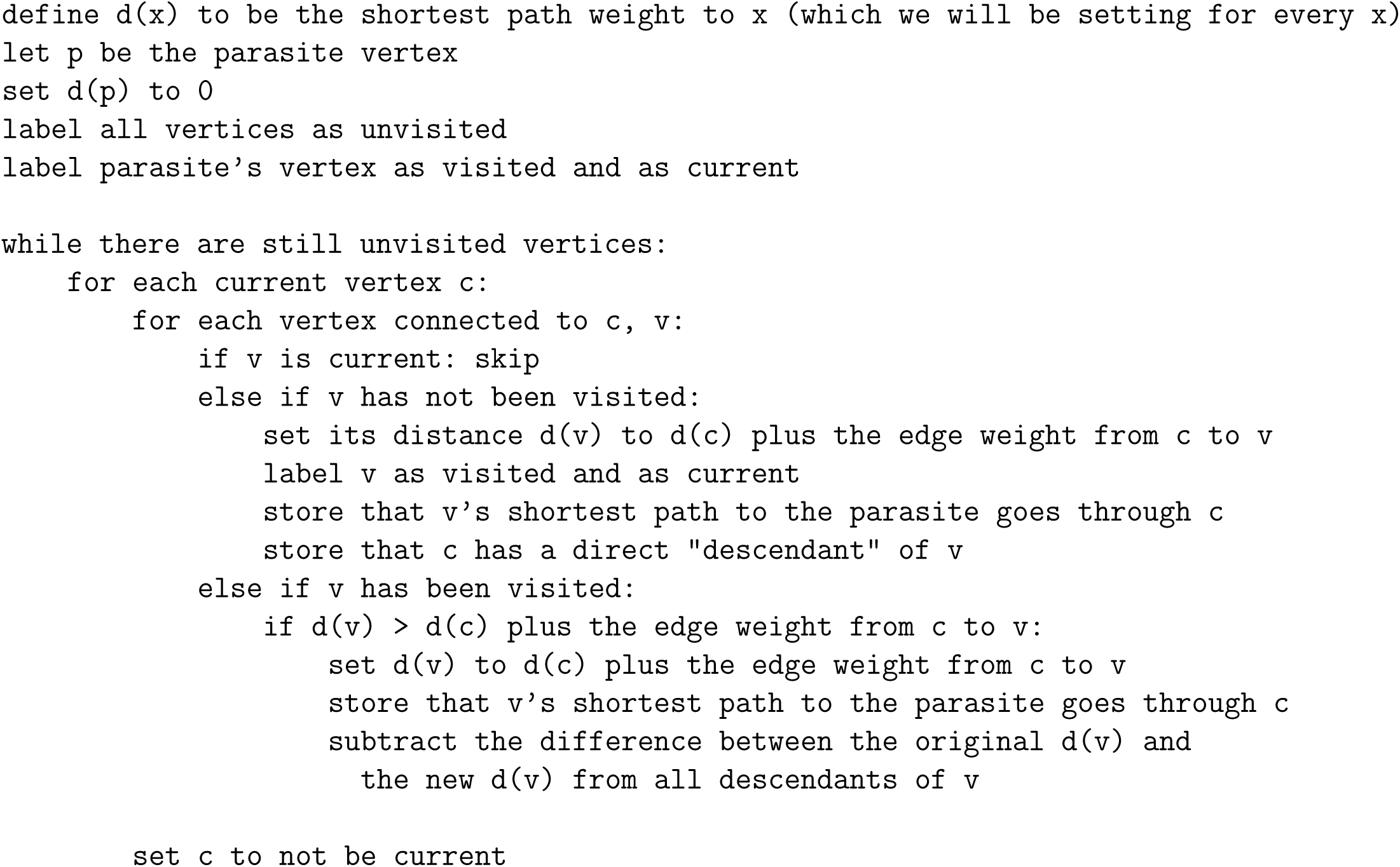

This algorithm is efficient for our problem of finding *every* shortest path in the graph. Other algorithms may be used for other problems and other graphs.

### Positions in space

Due to imperfections in tracing cell positions or the structure, as well as potential variations in sinusoid diameter or the microscope’s visibility, especially near the boundaries of the imaging volume, cells’ recorded positions and the sinusoids’ vertices’ corresponding positions in space may not be close to each other. From the 10*µ*m diameter estimate for most sinusoids, we surmise that with perfect tracing, all recorded cell positions would be within 5*µ*m of a vertex. Due to challenges in imaging and tracings, this is not always the case (**Supplemental Figure S3**).

**Movie 1:**
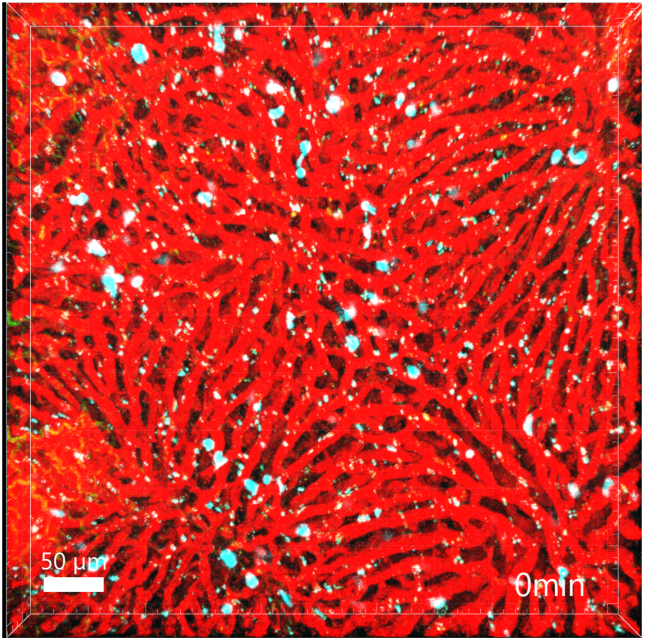
Imaging movements of liver-localized CD8 T cells within labeled liver sinusoids. We performed experiments as described in Figure 1 in 3 individual mice. In the movie (CTV5) red denotes rhodamine dextran-labeled liver sinusoids, and cyan are *Plasmodium*-specific liver-localized CD8 T cells (OT-I). The parasite is expressing GFP. Bar scale is 50 *µ*m. The volume of 512 *×* 512 *×* 48 *µ*m was scanned every 30 sec.

**Movie 2:**
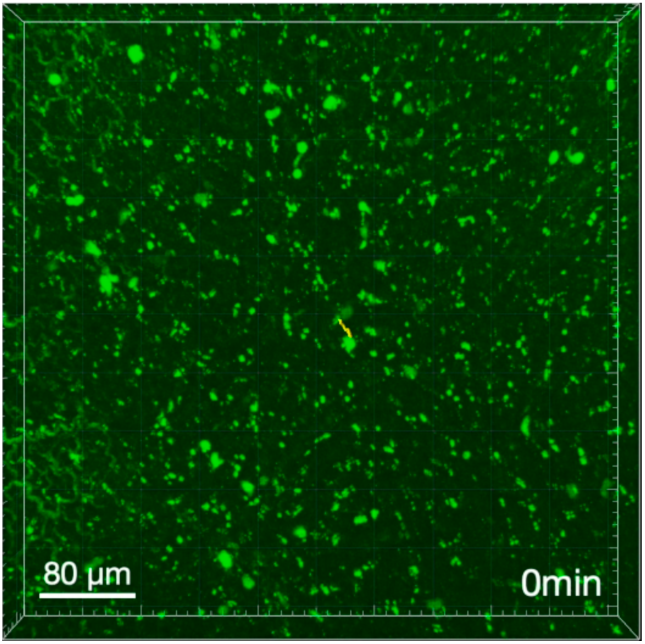
Tracking position of the Plasmodium sporozoites in the liver. We followed the location of the parasite identified during movie acquisition and tracked its position using the Spots function for GFP channel in Imaris. See Movie 1 for additional details.

**Movie 3:**
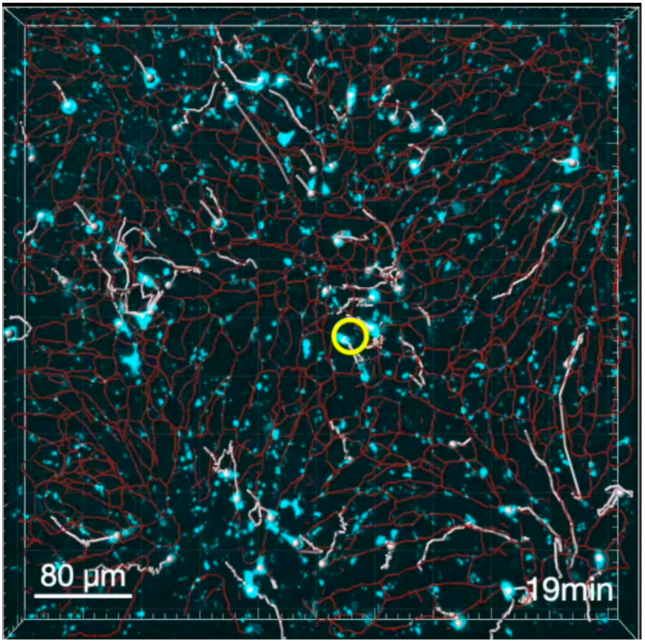
Tracking position of the liver-localized CD8 T cells. We used the Spots function in Imaris to track movements of liver-localized CD8 T cells over time. The yellow circle denotes the location of the Plasmodium liver stage. There was a tissue shift after 19 min of the movie that was accounted for by using the Reference Frame tool in Imaris (see Materials and methods and Movie 1 for additional details); for illustrative purposes this movie was cut short to avoid tissue shifts (and associated shifts in track positions).

**Movie 4:**
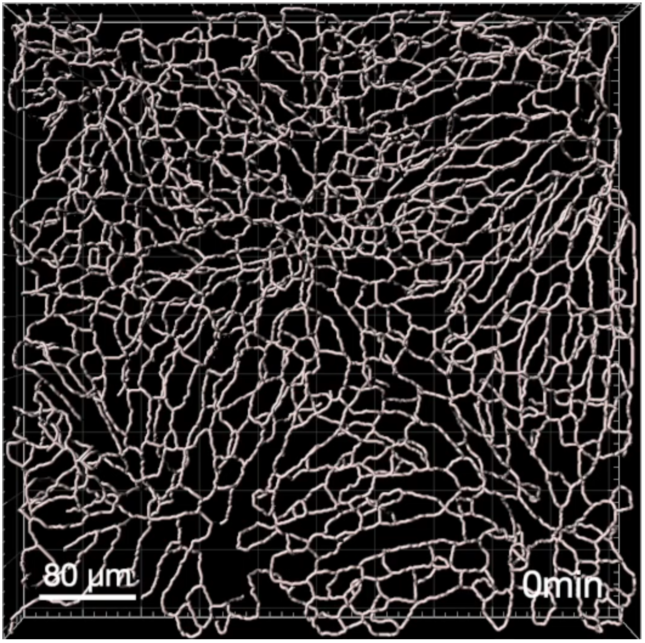
Tracking and quantifying liver sinusoids in Imaris. We injected rhodamine dextran intravenously and imaged the murine livers 15 min later. We then used the Filament tracer tool in Imaris to quantify liver sinusoids in the imaging volume. See Movie 1 for additional details.

**Movie 5:**
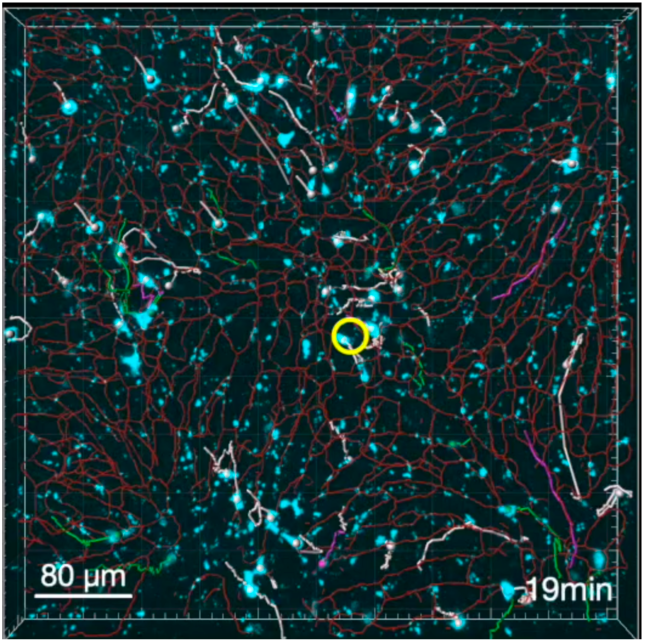
Movement of liver-localized CD8 T cells that are attracted or repulsed towards the liver stage via sinusoids. We used our graph-based metric to determine which T cells appeared to be attracted or repulsed towards the liver stage (see Materials and methods and Movie 1 for additional details).

**Supplemental Figure S1:**
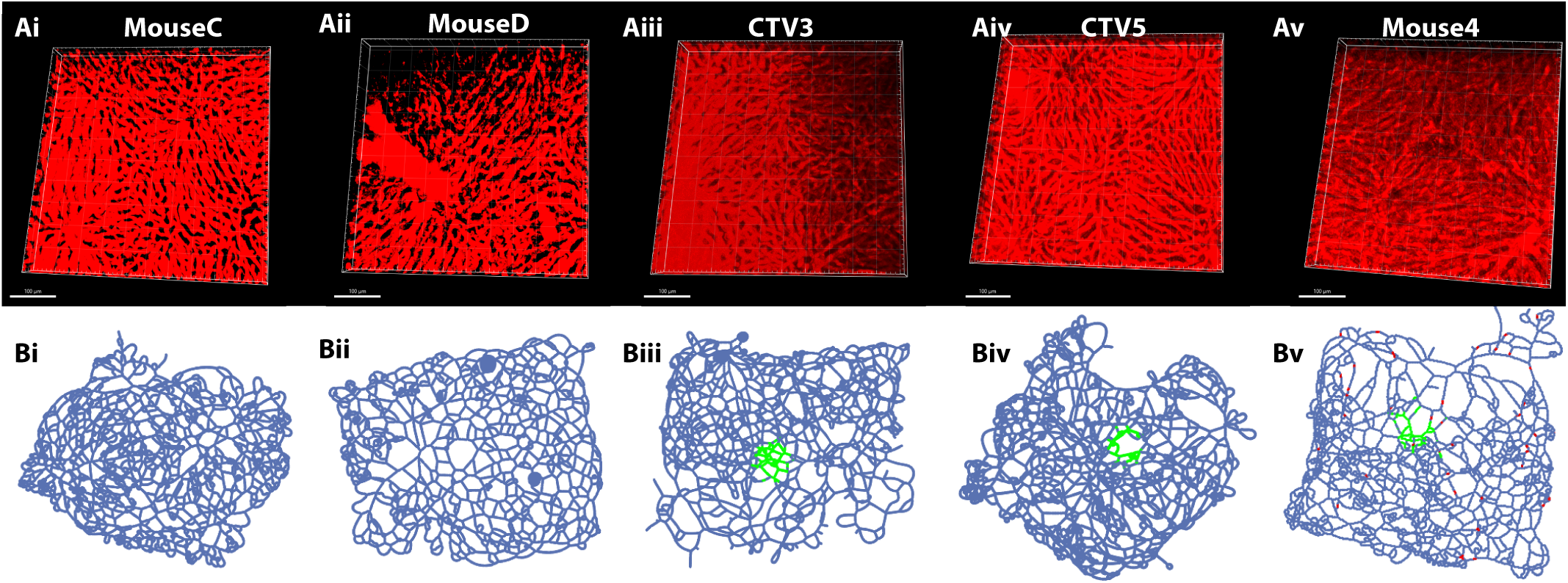
Rhodamine dextran-labeled liver sinusoids and corresponding graphs from five independent experiments. Out of five experiments (**Table 1**) two experiments (MouseC, **i** and MouseD, **ii**) did not involve infection, while the other three experiments (CTV3, **iii**, CTV5, **iv**, and Mouse4, **v**) featured a single sporozoite per imaging volume. We processed images (**A**) with Imaris and generated graphs (**B**). All vertices corresponding to positions in space estimated to be within the sporozoite-infected hepatocyte are highlighted in green.

**Supplemental Figure S2:**
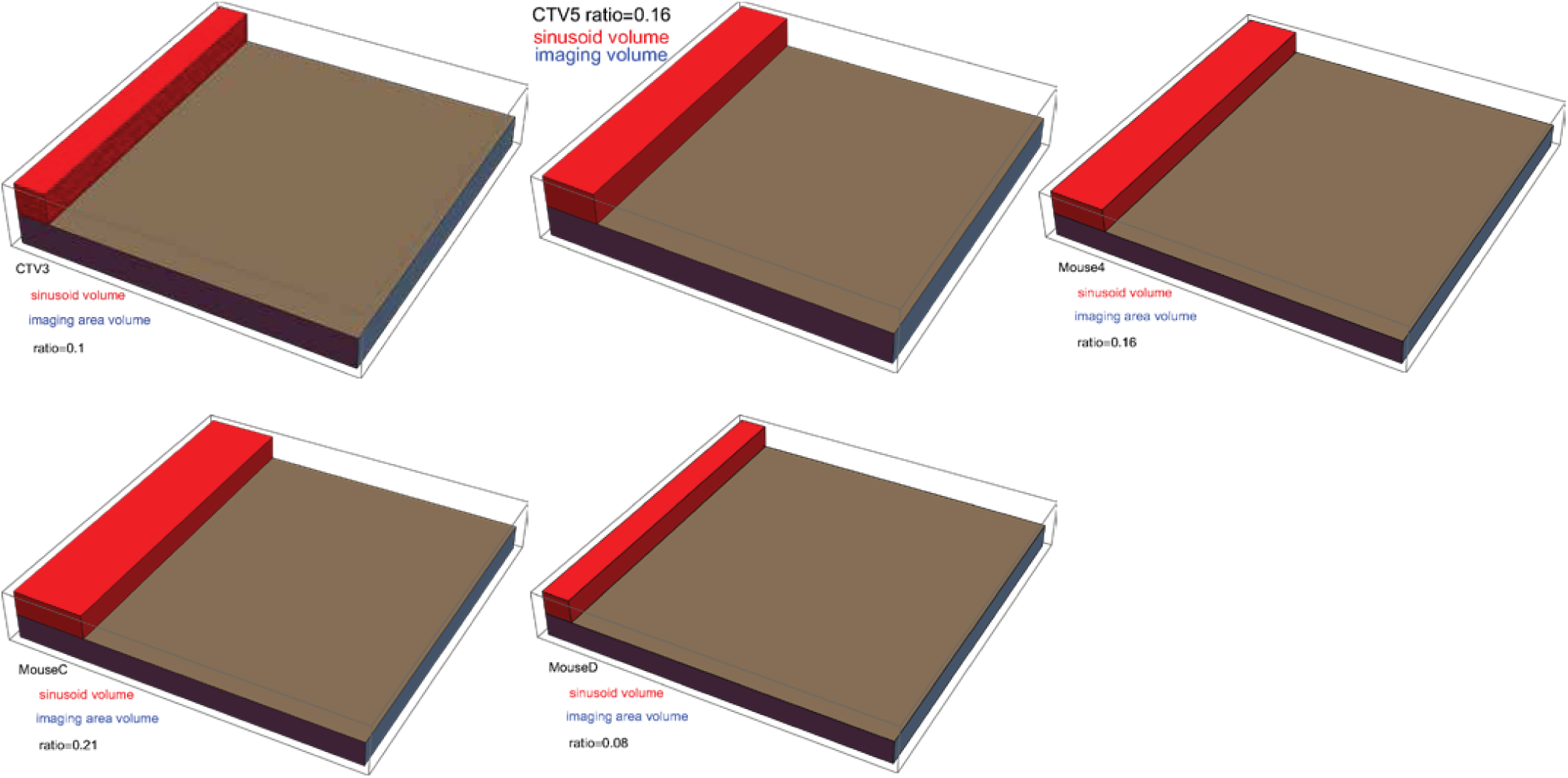
Accounting for the space available to T cells to move to liver sinusoids reduces the overall space by 80-90%. For each of the five movies, we calculated the total volume of the liver imaged (shown in brown) and the volume taken up by the sinusoids (shown in red).

**Supplemental Figure S3:**
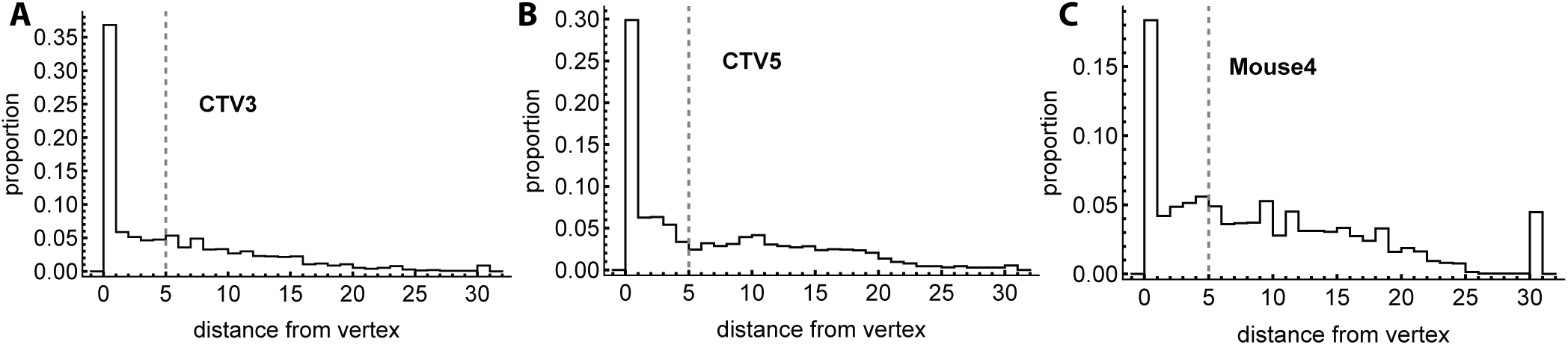
Most cells’ recorded positions are close to the sinusoid graph’s corresponding positions in space. Histograms show the proportion of cells’ visited positions in space that are within varied distances from the nearest vertex on the graph tracing. Sinusoids have a radius of about 5 *µ*m, so accounting for the possibility of the graph’s or the cells’ recorded positions being off-center, a perfect tracing would have no distance greater than 10 *µ*m.

**Supplemental Figure S4:**
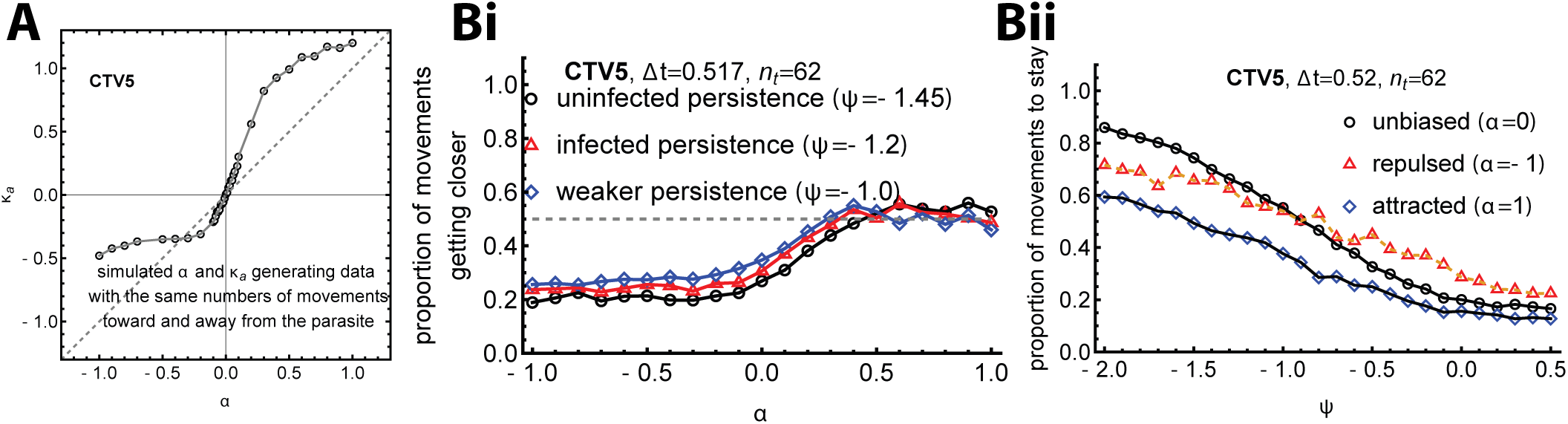
Impact of parameters *α* and *ψ* of graph-based simulations of T cell walks on inferred movement characteristics. We simulated movement of T cells on the graph from the CTV5 movie with varying degrees of attraction parameter *α* (A-Bi) or staying parameter *ψ* (Bii). **A** In a previous work, we simulated cells in open space using the VMF distribution, and estimated the proportion of cell movements to get closer to the parasite to be 1*/e^−κa^*^+1^, a function of the open space-based concentration parameter *κ_a_* [34]. A similar mathematical function may be impossible using our graph-based model, but we can find the proportions of movements closer/farther by pooling together movements of simulated cells with known values of *α*, shown in Figure 3A; we then match each *α* with the value of *κ_a_* that would generate similar proportions of movements closer/farther. **Bi** We estimate the proportion of movements getting closer to the parasite (a decreasing SPW) for varying values of *α* and *ψ*. **Bii** We estimate the proportion of movements staying in place for varying values of *ψ* and *α*.

**Supplemental Figure S5:**
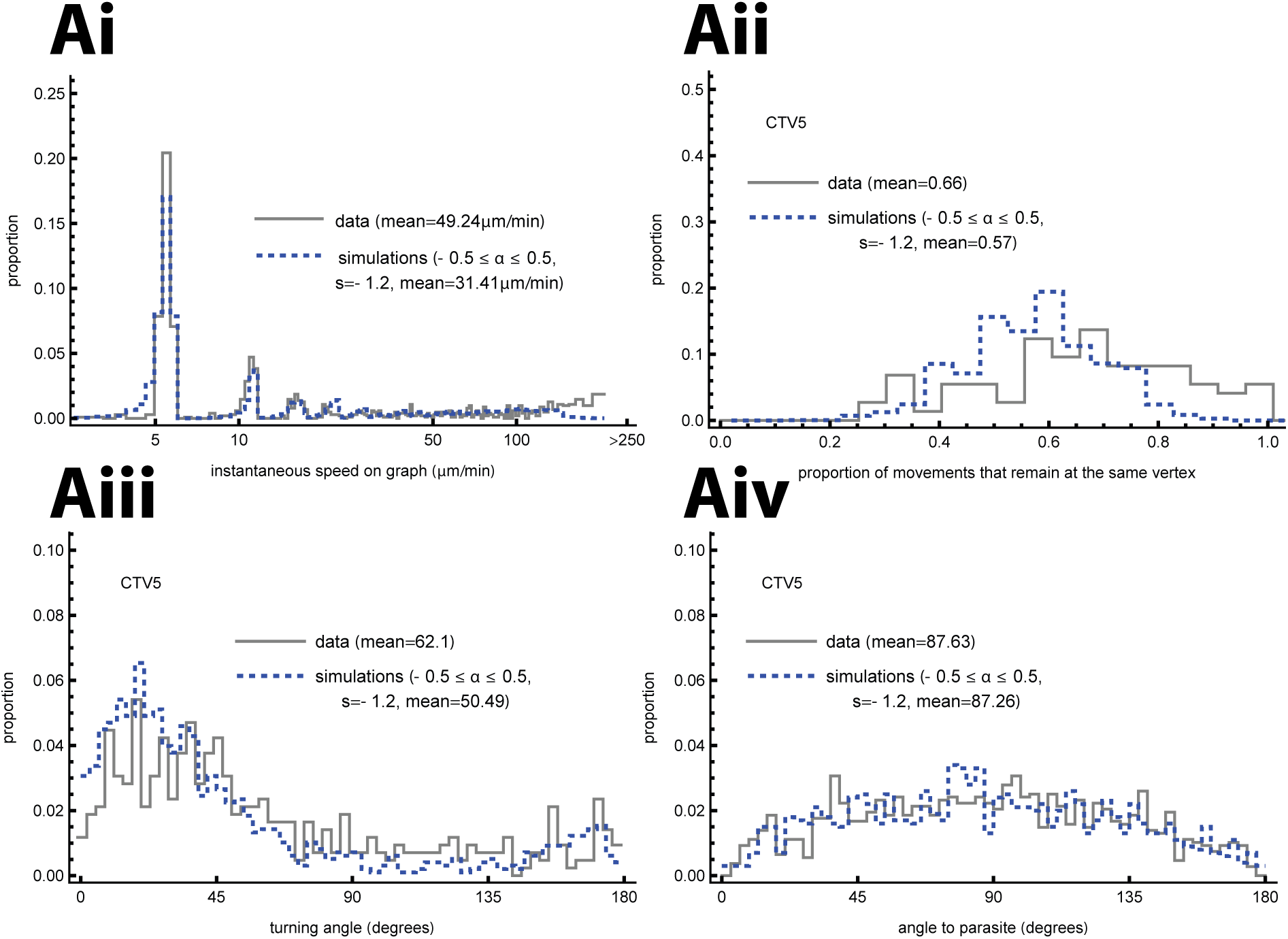
Simulated cells display similar movement characteristics as CD8 T cells observed experimentally. We analyzed T cell position data from the CTV5 experiment, simulated T cell movements on the graph from the CTV5 movie and compared basic movement characteristics of actual and simulated cells on the graph. Characteristics of actual T cells are shown with black solid lines, while characteristics for simulated cells are shown with blue dashed lines. **Ai**: We compare speeds of cells moving on the graph. **Aii**: We compare proportions of movements at which the cell stays in place. **Aiii**: We compare turning angles, i.e. the angle between subsequent vectors of movement of a cell. **Aiv**: We compare angles to the parasite, i.e. the angle between a cell’s movement vector and a vector from the cell’s position to the parasite. Parameters used in simulations are *α* = 0 and *ψ* = *−*1.2. Note that the average cell speed in Ai is 16.9 *µ*m*/*min, much lower than the average instantaneous speed.

**Supplemental Figure S6:**
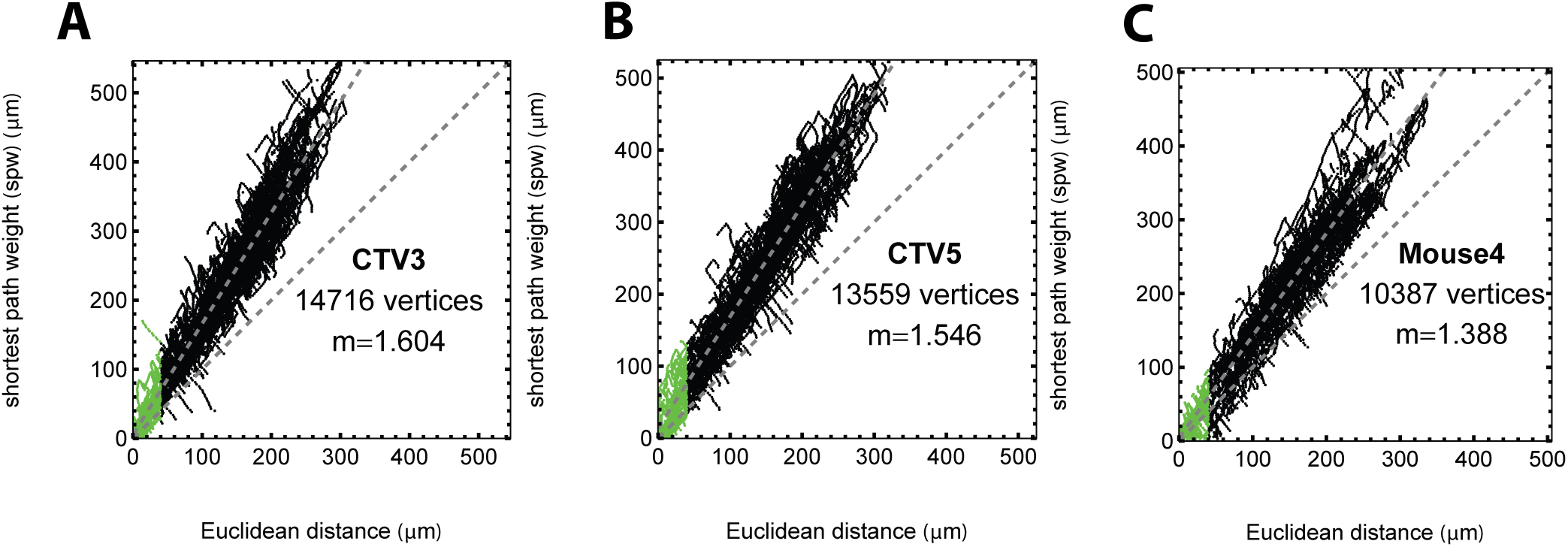
The actual distance between a T cell and the infection is longer than the Euclidean distance. For every position of every T cell, we calculated the Euclidean (3D) distance to the parasite and the actual distance to the nearest “infected” vertex on the graph (shortest path weight) for three experiments (CTV3, CTV5, and Mouse 4 shown in **A-C**, respectively). The slope *m* denotes the regression slope between the two metrics. Points in green show vertices corresponding to the parasite.

**Supplemental Figure S7:**
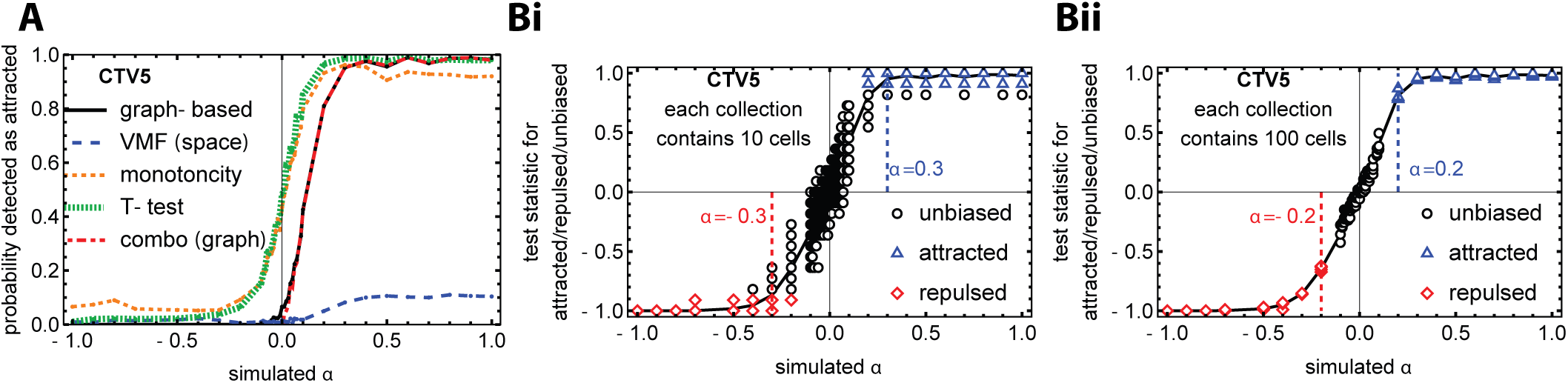
The novel graph-based test is most accurate at detecting cells as unbiased, attracted or repulsed. We simulated T cell movement for 300 steps with repulsion (*α <* 0), attraction (*α >* 0) or as unbiased (*α* = 0) on a graph from the CTV5 experiment and tested how methods based on different metrics are able to accurately detect cells as attracted, repulsed, or unbiased (see Materials and methods for detail). **A**: We test each simulated cell using our primary tests (the graph-based and space-based VMF) and alternate graph-based tests (monotonicity and T test), all of which test per cell, as well as the graph-based “combo” test on graph-based results from collections of 10 cells. **B**: We approximate the minimum values of *α* at which the combo test consistently detects attraction or repulsion for a collection of 10 cells (**i**) or 100 cells (**ii**).

**Supplemental Figure S8:**
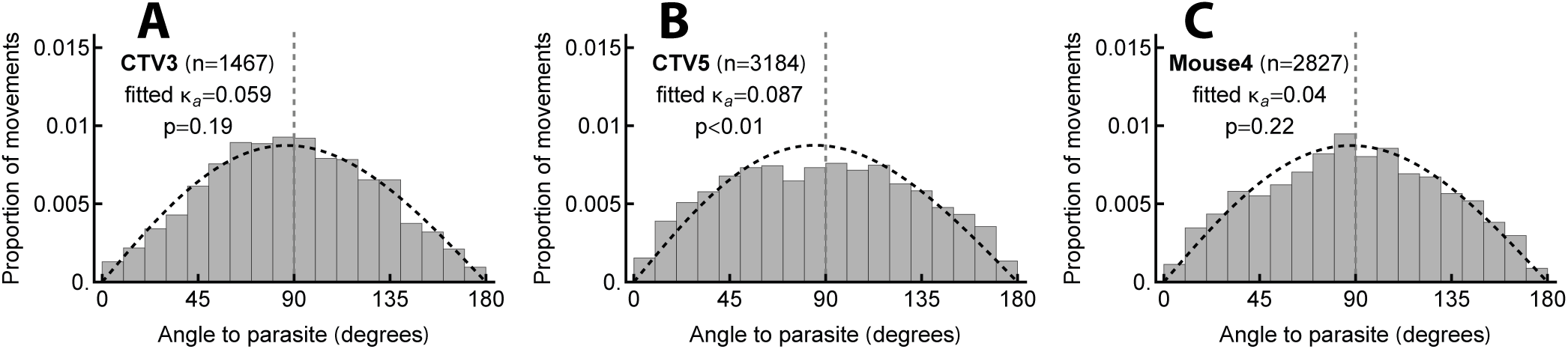
Open space/3D metric based on the VMF distribution detects a small bias in moving T cells when there is a small cluster around the liver stage. For three experimental movies (CTV3 in **A**, CTV5 in **B**, and Mouse4 in **C**) we calculated the angle to infection (angle between the movement vector of a T cell and the vector to the parasite) for every movement of every CD8 T cell, estimated the concentration parameter *κ_a_* of the VMF distribution using the likelihood method (**eqn. (3)**), and evaluated its deviation from the null model that has *κ_a_* = 0.

**Supplemental Figure S9:**
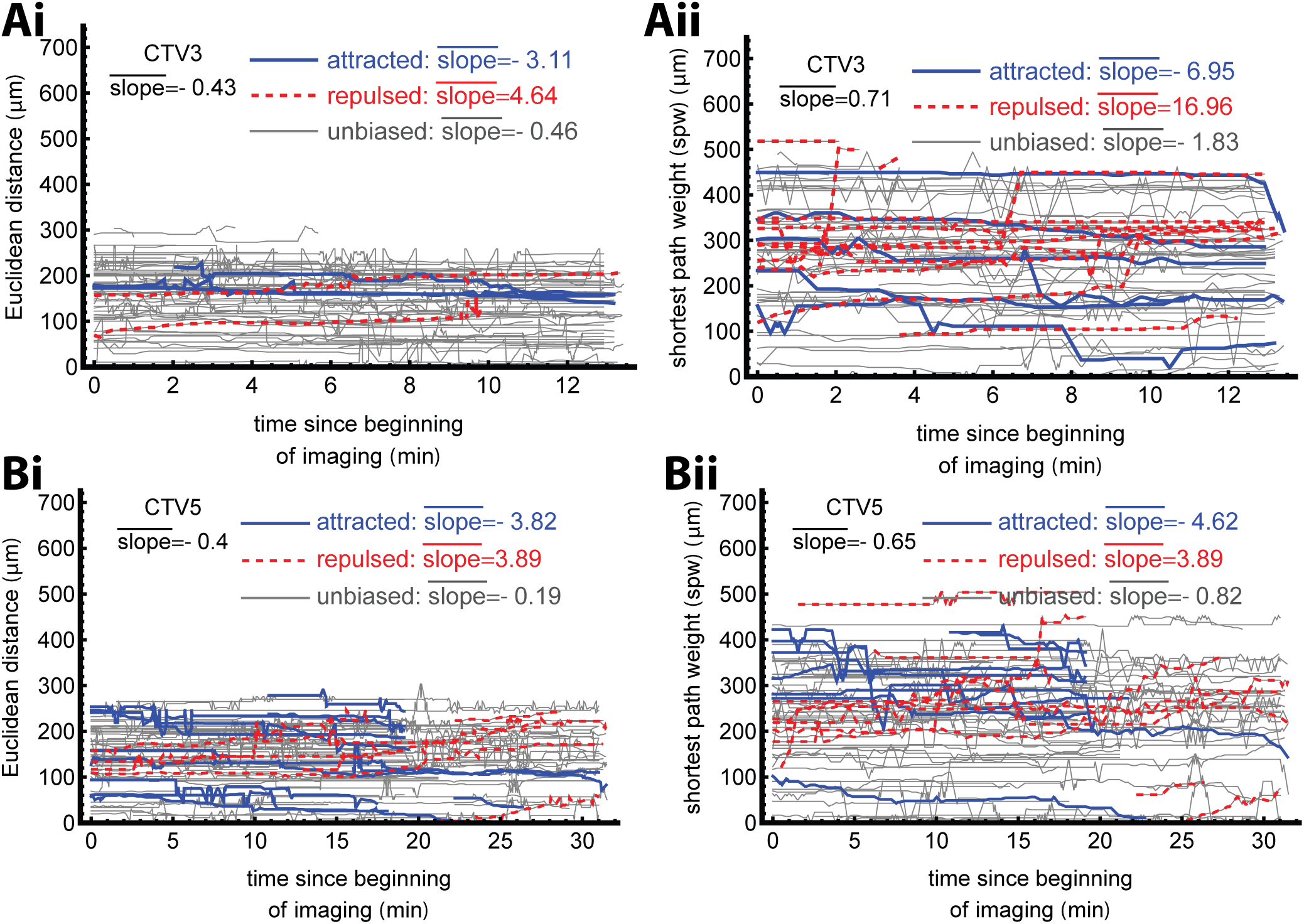
Cells detected as attracted move closer to the parasite. We estimated change in Euclidean distance (**Ai**/**Bi**) or SPW (**Aii**/**Bii**) for T cells in CTV3 (no cluster, **A**) and CTV5 (small clustered, **B**) datasets. **i**: The Euclidean distances of each cell are plotted over time, with lines of cells detected significantly as attracted or repulsed using the space-based VMF distribution-based test (**eqn. (3)**) highlighted in blue and red respectively. **ii**: The SPWs of each cell are plotted over time, with lines of cells detected significantly as attracted or repulsed using the graph-based test (**eqn. (6)**) highlighted in blue and red respectively.

**Supplemental Figure S10:**
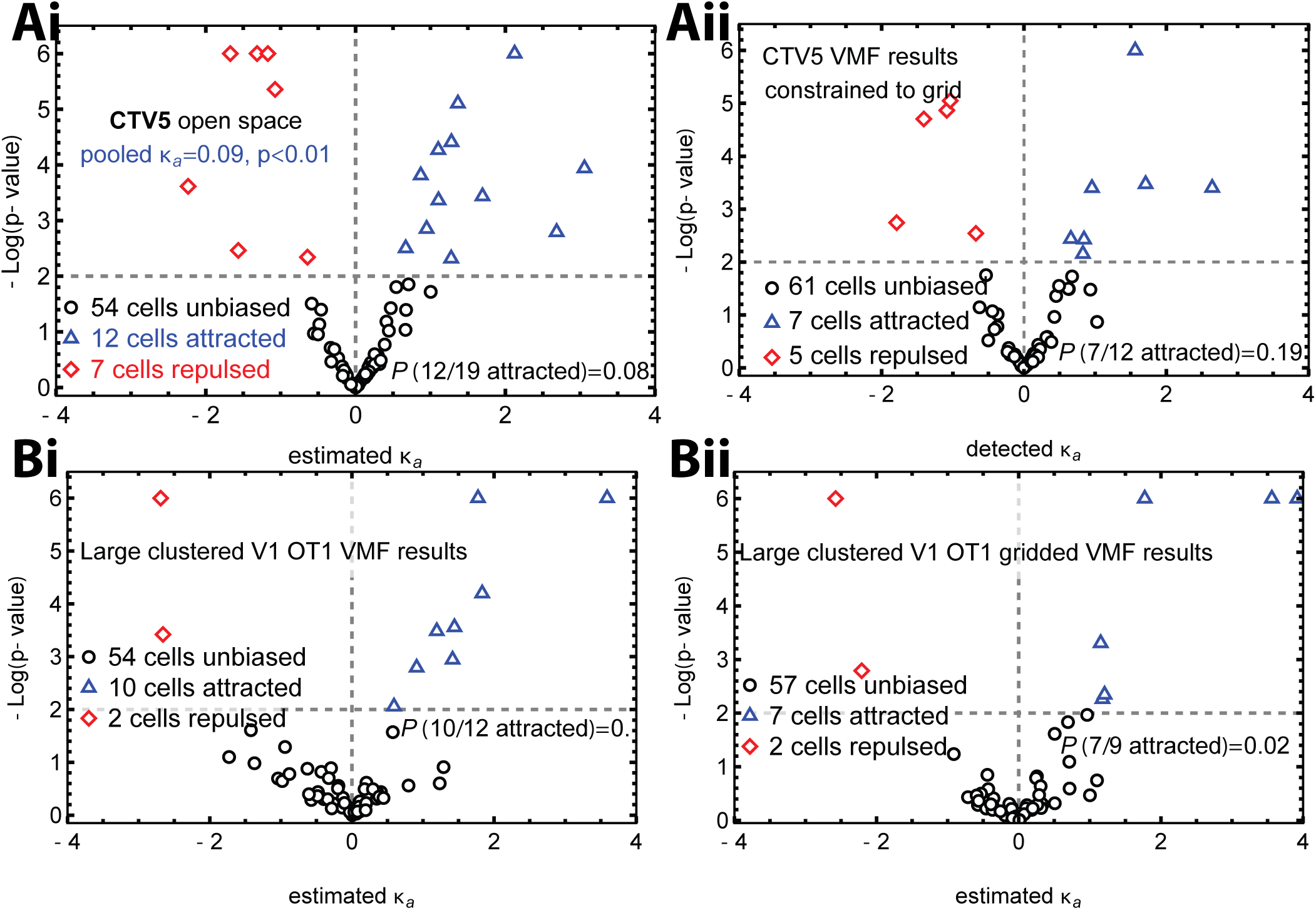
Minimizing small movements of cells without taking the structure into account does not change the conclusions based on the open space/3D metric. The format of the panels is as described in Figure 4, plotting p-values against detected values of *κ_a_* for each cell in a dataset. **Ai**: Equivalent to Figure 4Di, these are results for the small clustered dataset. **Aii**: We use angles to the parasite between the positions in space matching the nearest positions on a grid with side length 3*µ*m, *not* taking into account the true constrained structure but nevertheless removing small movements of cells which may be artifacts of imaging, floating in the blood, shifting of the mouse or microscope, or other factors unrelated to potentially biased movement. **Bi**: We use data from a previous paper, whose experiments were similar but did not image sinusoids; this dataset featured a “large cluster” of around 5-6 T cells close to the parasite and we detected attraction [34]. **Bii**: We map the previous paper’s cells’ positions to the nearest vertices on a 3D grid with side length 3*µ*m, as in **Aii**.

**Supplemental Figure S11:**
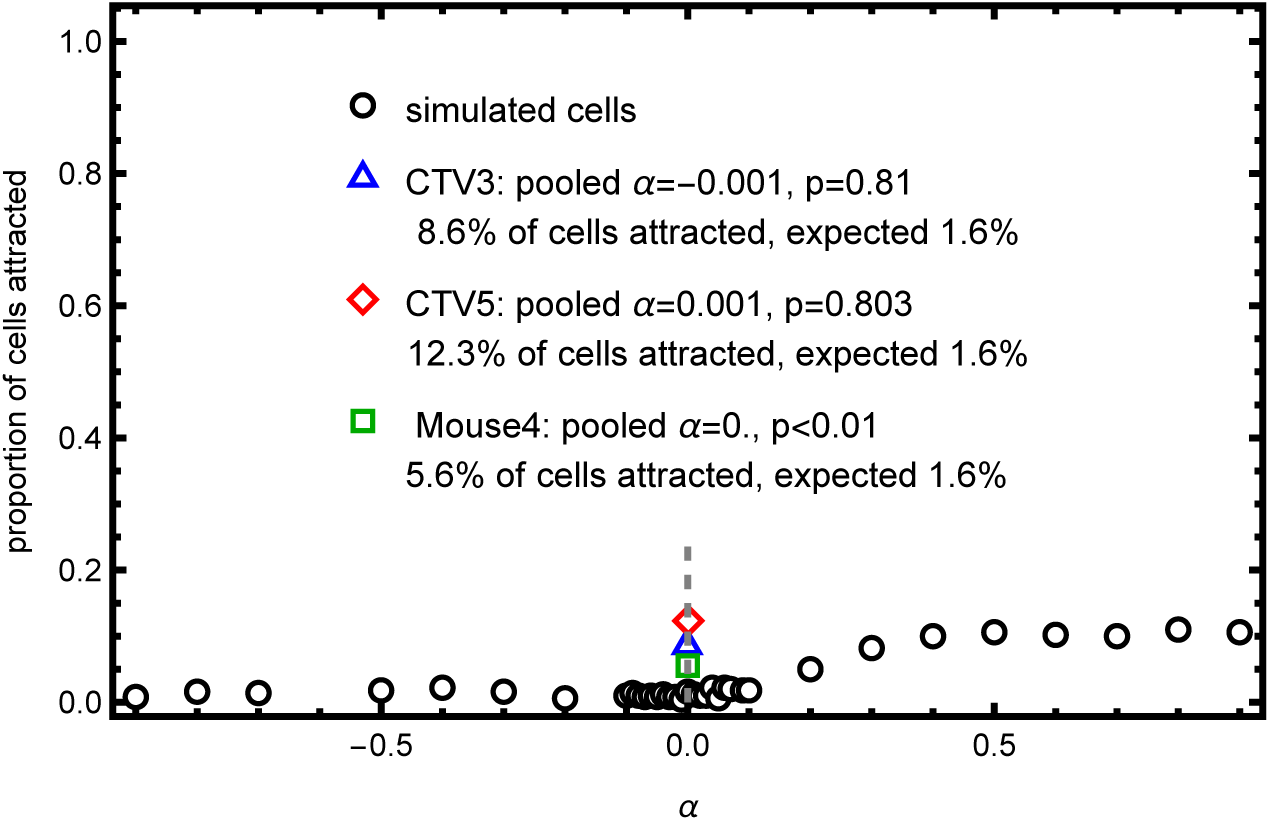
With the graph-based metric, while T cells demonstrate no attraction a higher proportion of T cells is detected as biased than expected. For simulated cells with *α* between −1 and 1, we calculate the proportion of cells detected as attracted. For each real dataset, we also calculate that proportion, as well as the estimated *α* on pooled data from that dataset; we hypothesize that the “expected” proportion of cells detected as attracted for that dataset should be the value for simulated cells moving with that pooled value of *α*.

**Supplemental Figure S12:**
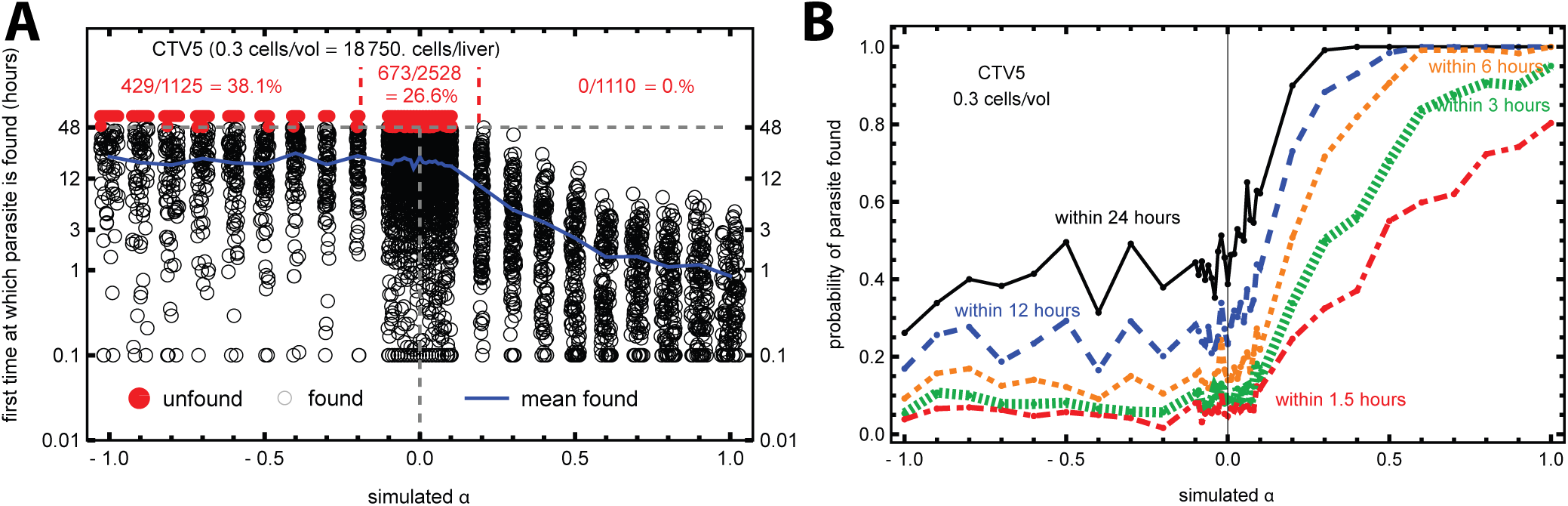
Even fewer than one cell per imaging volume moving randomly without attraction is able to effectively locate the infection site. We simulated movement of T cells assuming that one T cell appears only 1/3 of the time in the imaging volume using data from experiment CTV5 (see Figure 5 for detail). Having 1/3 cells per imaging volume corresponds to 18,750 T cells per liver. **A**: The time to find the infection for different values of *α* out of 100 simulations. Red points indicate runs for which the parasite was not found within 48 hours. The dashed blue line indicates the average time at which the parasite is found for different values of *α*. **B**: The probability that T cells find the parasite within different times after infection as the function of *α*.

**Supplemental Figure S13:**
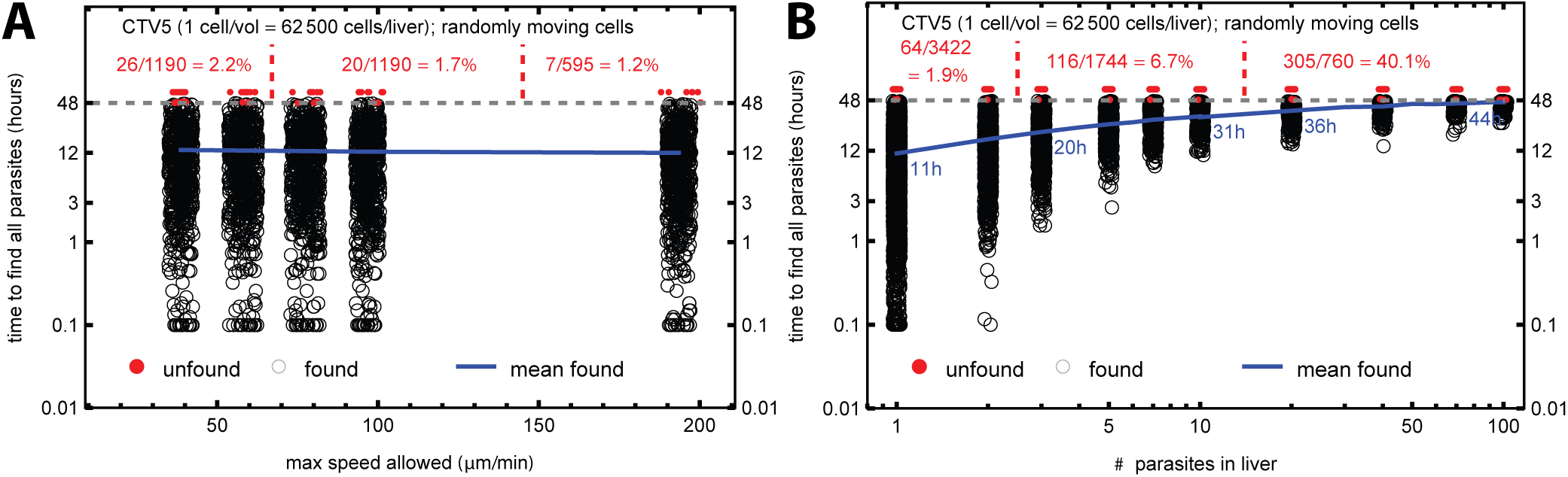
Reducing the maximum speed of T cells or searching for several parasites does not impact the time to find the infection on graphs. We simulated a search by T cells for a single parasite while restricting the maximum speed T cells may have (**A**) or simulating a search for multiple parasites, assuming that the parasites are located in different areas of the liver and are thus independent (**B**). Notations in panels are similar to Figure 5A.

**Supplemental Figure S14:**
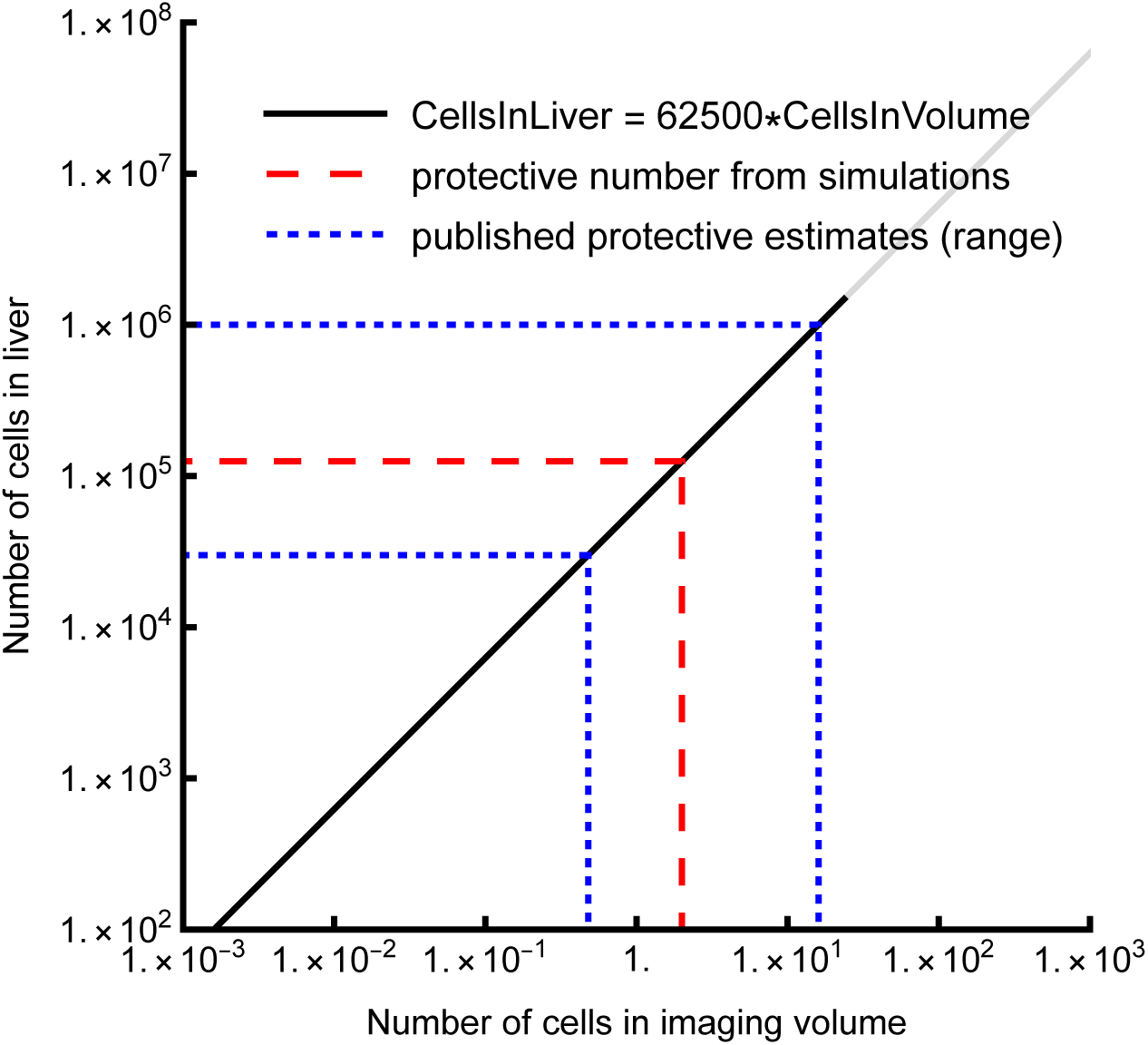
The number of T cells sufficient to protect from infection, predicted by simulations of unbiased T cells, is within the experimentally observed range of protective liver-localized T cells. We plot the rate of previously published experimentally-derived numbers of liver-localized CD8 T cells sufficient to protect against exposure to Plasmodium sporozoites (dashed blue lines, [32]) and the protective number of T cells found in our simulations (dashed red lines, see Figure 5C).

